# Highly clade-specific biosynthesis of rhamnose: present in all plants and in only 42% of prokaryotes. Only *Pseudomonas* uses both D- and L-rhamnose

**DOI:** 10.1101/854612

**Authors:** Toshi Mishra, Petety V. Balaji

## Abstract

Rhamnose is a constituent of lipo- and capsular polysaccharides, and cell surface glycoproteins. L-rhamnose is biosynthesized by the rml or udp pathway and D-rhamnose by the gdp pathway. Disruption of its biosynthesis affects survival, colonisation, etc. Rhamnosides are commercially important in pharmaceutical and cosmetics industries. HMM profiles were used to investigate the prevalence of the three pathways in completely sequenced genomes and metagenomes. The three pathways are mutually exclusive except in *Pseudomonas* which has both rml and gdp pathways. The rml pathway is restricted to bacteria (42% genomes), archaea (21%) and bacteriophages, and absent in eukaryotes and other viruses. The gdp pathway is restricted to *Pseudomonas* and *Aneurinibacillus*. The udp pathway is primarily found in plants, fungi and algae, and in human faecal metagenomic samples. The rml pathway is found in >40% genomes of Actinobacteria, Bacteroidetes, Crenarchaeota, Cyanobacteria, Fusobacteria and Proteobacteria but in <20% genomes of Chlamydiae, Euryarchaeota and Tenericutes. The udp pathway is found in all genomes of Streptophyta, <=25% genomes of Ascomycota and Chordata, and none of the genomes of Arthropoda and Basidiomycota. Some genera which lack any of these pathways are *Chlamydia*, *Helicobacter*, *Listeria*, *Mycoplasma*, *Pasteurella*, *Rickettsia* and *Staphylococcus*. Organisms such as *E. coli* and *Salmonella enterica* showed significant strain-specific differences in the presence/absence of rhamnose pathways. Identification of rhamnose biosynthesis genes facilitates profiling their expression pattern, and in turn, better understanding the physiological role of rhamnose. Knowledge of phylogenetic distribution of biosynthesis pathways helps in fine graining the taxonomic profiling of metagenomes.

**AUTHOR SUMMARY:** In the present study, we have investigated the prevalence of rhamnose biosynthesis pathways in completely sequenced genomes and metagenomes. It is observed that the prevalence of rhamnose is highly clade specific: present in all plants but in less than half of all prokaryotes. Among chordates, only the Chinese rufous horseshoe bat has rhamnose biosynthesis pathway and this exclusive presence is quite baffling. The effect of disrupting rhamnose biosynthesis has been reported in a few prokaryotes and all these cases pointed to the essentiality of rhamnose for critical physiological processes such as survival, colonisation, etc. In this background, it is surprising that many of the prokaryotes such as *Escherichia coli* and *Salmonella enterica* show significant strain-specific differences in the presence/absence of rhamnose pathway. This study will facilitate the experimental characterization of rhamnose biosynthesis genes in organisms where this pathway has not been characterised yet, eventually leading to the elucidation of the biological role of rhamnose. Phylum-, genus-, species- and strain-level differences found with respect to presence of rhamnose biosynthesis pathway genes can be used as a tool for taxonomic profiling of metagenome samples. This study could also annotate a significant number of orphan proteins in the TrEMBL database.

## INTRODUCTION

Rhamnose is 6-deoxy-mannose. Both L- and D-enantiomers of rhamnose are found in nature. Its presence has so far been reported in bacteria, archaea, plants and fungi. It is found as part of polysaccharides (1), glycoproteins (2) and small molecules such as flavonoids, terpenoids and saponins (3–5). L-rhamnose is a common component of the O-antigen of lipopolysaccharides (LPS) of Gram-positive bacteria such as *Lactococcus lactis* (6) and *Enterococcus faecalis* (7, 8), and Gram-negative bacteria such as *Salmonella enterica* (9), *Shigella flexneri* (10) and *Escherichia coli* (11). D-Rhamnose is a constituent of LPS of Gram-negative bacteria such as *Pseudomonas aeruginosa* (12) and *Pseudomonas syringae* (13, 14), and Gram-positive thermophilic bacterium *Aneurinibacillus thermoaerophilus* (15). The structure of rhamnose-containing glycan moiety has been elucidated in some bacteria. For e.g., the 93-kDa S-layer glycoprotein SgsE of *Geobacillus stearothermophilus* NRS 2004/3a is O-glycosylated at Ser/Thr residues (2). In this, rhamnan chains consisting of 12-18 repeating units of [-L-Rha-α1,3-L-Rha-β1,2-L-Rha-α1,2-] are linked to Ser/Thr through the core disaccharide [L-Rha-α1,3-L-Rha-α1,3-]. Occasional 2-O-Me modification of terminal residues and inclusion of a third rhamnose in the core bring about microheterogeneity.

L-rhamnose is a constituent of the glycan moiety of glycoproteins that make up flagella, fimbriae and pili. For e.g., flagellin from *P. syringae* pv. tabaci 6605 consisted solely of L-Rha whereas the *P. syringae* pv. glycinea race 4 flagellin contained L-Rha and D-Rha in ∼4:1 ratio (14). Fap1 adhesin is a major fimbrial subunit of *Streptococcus parasanguis* and its glycan contains rhamnose, glucose, galactose, GlcNAc and GalNAc in the ratio of 1:29:5:39:1 (16). Glycosylation of flagellin has been shown to be essential for bacterial virulence and host specificity (17, 18).

Three major pectic polysaccharides (homogalacturonan and rhamnogalacturonans RG-I and RG-II) are present in cell walls of plants. Pectic polysaccharides consist of [-L-Rha-α1,4-D-GalA-α1,4-] repeating units and play a major role in the development and growth of all vascular plants (19). Rhamnosides exhibit a wide range of biological activities like anti-inflammatory (20, 21), anti-viral (22, 23), anti-oxidant (24, 25) and anti-cancer activities (26–28).

Three pathways are known for the biosynthesis of rhamnose and these will henceforth be referred to as rml, gdp and udp pathways for brevity (Figure 1). (d)TDP-L-rhamnose, synthesised via the rml pathway, has been reported in bacteria and archaea. *Pseudomonas* additionally uses GDP-D-rhamnose synthesised via the gdp pathway. Plants and fungi use UDP-L-rhamnose biosynthesised via the udp pathway. This pathway has only two steps since Uger has two activities viz., 3,5-epimerase and 4-reductase in a single active site. In some plants such as *Arabidopsis thaliana*, the pathway has just one step since a fusion protein viz., RHM consists of both Ugd and Uger activities (29).

**Fig. 1.**
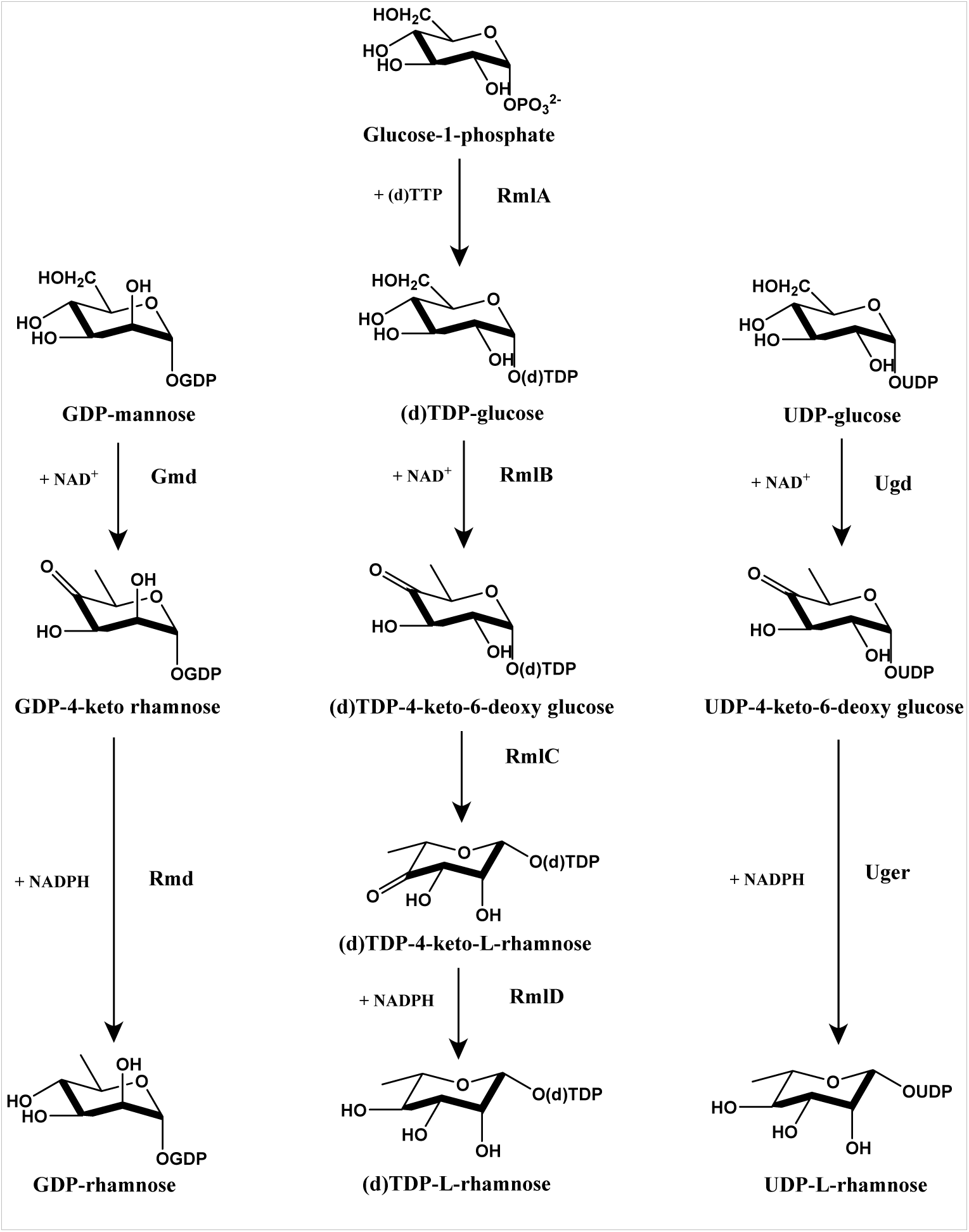
Pathways for the biosynthesis of nucleotide-rhamnose. The gdp pathway (*left column*) is for the biosynthesis of GDP-rhamnose whereas the rml (*middle column*) and udp (*right column*) pathways are for the biosynthesis of (d)TDP-L-rhamnose and UDP-L-rhamnose, respectively. For brevity, these three pathways are referred to as the rml, gdp and udp pathways. UDP-glucose and GDP-mannose (starting substrates for udp and gdp pathways) are also biosynthesized from glucose-1-phosphate, but these segments of the pathway are not unique to the rhamnose biosynthesis pathways. Enzyme activities are as follows: RmlA, Glucose-1-phosphate thymidylyltransferase; RmlB, (d)TDP-glucose 4,6-dehydratase (retaining); RmlC, (d)TDP-4-keto-6-deoxy-glucose 3,5-epimerase; RmlD, (d)TDP-4-keto-L-rhamnose reductase; Ugd, UDP-glucose 4,6-dehydratase (retaining); Uger, UDP-4-keto-6-deoxy-glucose 3,5-epimerase, 4-reductase; Gmd, GDP-glucose 4,6-dehydratase (retaining); Rmd, GDP-4-keto-rhamnose reductase.

Bacterial cell surface polysaccharides contain rhamnose and play important roles in cell growth and development, survival and interaction between bacteria. Disruption of the rhamnose biosynthesis pathway in *Enterococcus faecalis* attenuates the pathogen in a mouse model (30). Deletion or disruption of the rml pathway in *Pseudomonas aeruginosa* is effectively lethal (31). Deletion of *rmlB* or *rmlD* in *Vibrio cholerae* results in defective colonisation (32). Deletion of any of the four *rml* genes in *Streptococcus mutans* inhibits cell-wall polysaccharide synthesis and mutants cannot initiate or sustain an infection (33). In the uropathogenic strain O75:K5 of *E. coli*, lack of functional RmlD leads to loss of serum resistance (34). In *Mycobacterium tuberculosis*, L-rhamnose covalently links arabinogalactan to peptidoglycan layer and inhibitors of rml pathway enzymes result in growth inhibition (35–37). These observations confirm the role of rhamnose in bacterial pathogenicity. To date, neither rhamnose nor the genes responsible for its synthesis have been identified in humans. Thus, inhibitors of rhamnose biosynthesis pathways can be used for novel therapeutic intervention.

Elucidation of the pathways of glycosylation and biosynthesis of glycan building blocks can alleviate the challenges associated in relating structure of glycans to their functions. It will also greatly facilitate chemoenzymatic synthesis of building blocks. In addition, knowledge of the pathways will help in investigating the regulation of their expression. In this background, the present study was undertaken to identify rhamnose biosynthesis pathways in sequenced genomes. Protein Data Bank and Swiss-Prot database were mined to extract protein family-specific sequence and/or structural patterns. Sequence patterns in the form of hidden Markov model (HMM) profiles were used to identify rhamnose biosynthesis genes. Genus-, species- and strain-specific differences in the use of rhamnose were also analysed. Genomic context was used to assign the biological process level annotations in prokaryotes. Human-associated and environmental metagenomes were analysed.

## RESULTS

### Specificity and sensitivity of HMM profiles

Ascertaining the specificity or sensitivity of an HMM profile is not straightforward since the only reliable validation is by direct enzyme activity assay. This is impractical because the number of proteins to be assayed is extremely large; in fact, HMMs are typically used to narrow down the possible activities a protein may be associated with. In the background of this caveat, the performance of the HMM profiles generated in this study was assessed using proteins which have molecular function annotations in the Swiss-Prot, Pfam and CATH databases.

#### Assessment using fusion proteins

Same protein was obtained as hit for both RmlC and RmlD in 69 of the 21,964 genomes. Lengths of these hits (441-494 residues) suggested that these might be fusion proteins. Inspection of alignments showed that distinct regions of the proteins align to RmlC and RmlD profiles strengthening the possibility that they are indeed fusion proteins. This was taken advantage of to validate the bit score thresholds of these two profiles. Hits of RmlC and RmlD from the genome were paired together, irrespective of whether they form fusion proteins or not, and their scores against respective profiles were plotted against each other (Figure S1). It is seen that most of the RmlC-RmlD fusion proteins scored just above the threshold. Since only functionally interacting proteins are found as fused proteins, the choice of bit score threshold was taken as validated. The lower bit scores are suggestive of sequence divergence.

#### Assessment using genomic context

Co-occurrence of genes in a genome is a strong indicator of functional similarity at the level of biological process (38). Prokaryotic genomes which have homologs for all enzymes of pathway were chosen and homologs in such genomes were classified as contiguous, neighbourhood and dispersed. The distribution of bit scores of hits of these three categories was very similar (Figure S2) indicating the validity of bit score thresholds.

#### Assessment using the Swiss-Prot database

This database was chosen since annotation of entries in this database are manually curated. Several of the Swiss-Prot entries are already part of the Extend dataset (Swiss-Prot Validation sheet of rhamnose.xlsx) and expectedly, these are obtained as hits for all profiles. After excluding these, proteins which are annotated at molecular function level were found as hits only for RmlA, RmlC and RmlD profiles and all of these, except one, are false positives. For the RmlA profile, six GlmUs and one glucose-1-phosphate uridylyltransferase are obtained as hits. Some of the functionally important residues are not conserved in these 7 proteins: Q26, L88, N111 and V172 in GlmUs and Q26 and W223 in the uridylyltransferase (Q9HU22 numbering). An uncharacterized protein (Q9HDU4) is obtained as a hit for RmlB+Ugd profile. All the functionally important residues are conserved in this protein and this will be a true positive. For the RmlC profile, the protein Q9RR30 annotated as Probable dTDP-4-oxo-2,6-dideoxy-D-glucose 3,5-epimerase is obtained as a hit but catalytic residues are not conserved in this protein suggesting that this is unlikely to have catalytic activity. For the RmlD profile, Probable dTDP-4,6-dihydroxy-2-methyloxan-3-one 4-ketoreductase (Q9RR27) [involved in the biosynthesis of dTDP-L-olivose; Aguirrezabalaga et al., 2000] and Spore coat polysaccharide biosynthesis protein SpsK (P39631) are also obtained as hits. Both of these are false positives, since some of the functionally important residues are not conserved. There were no false negatives for any of the profiles.

#### Assessment using the Pfam database

Pfam is a database of protein families wherein proteins are grouped based on sequence / structure similarity or family-specific HMM profiles (40). Pfam families which include rhamnose biosynthesis pathway enzymes were identified by keyword search (primary protein name and gene name; Table 1): these are PF00483, PF00908, PF01370, PF04321 and PF16363 (Table S1). These five Pfam families contain proteins with a variety of molecular functions. Full-length sequences for all members of each these Pfam families were scanned using the corresponding HMM profiles generated in this study (Table 1 and Pfam Validation sheet of rhamnose.xlsx). It is found that all the profiles generated in this study have high specificity and some also have high sensitivity; detailed analysis are given in Table S1. Some of the entries from Pfam families PF01370 and PF16363 are annotated as Gmd but they are not hits for the Gmd profile; hence, these are likely to be RmlBs.

**Table 1.**
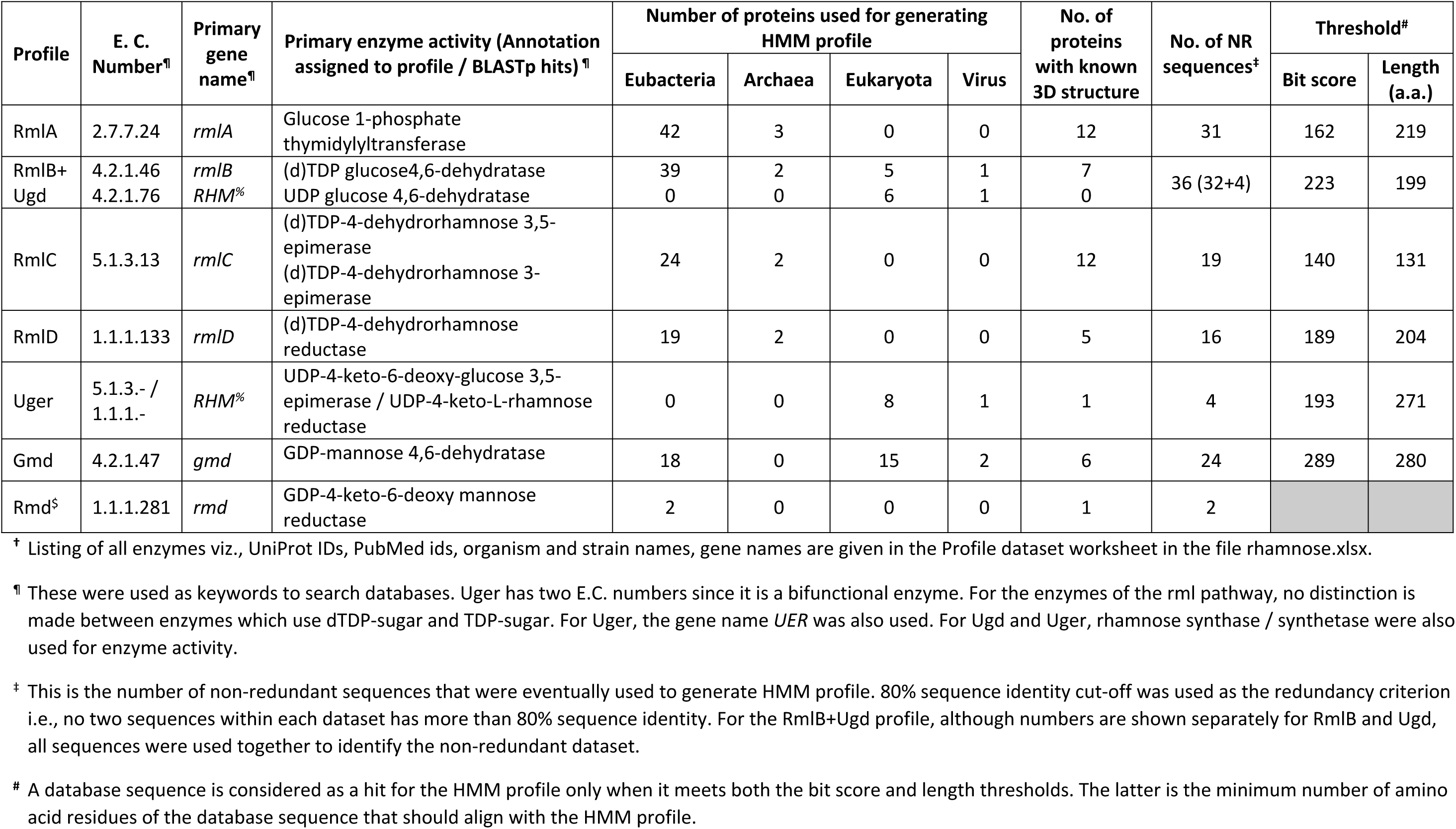
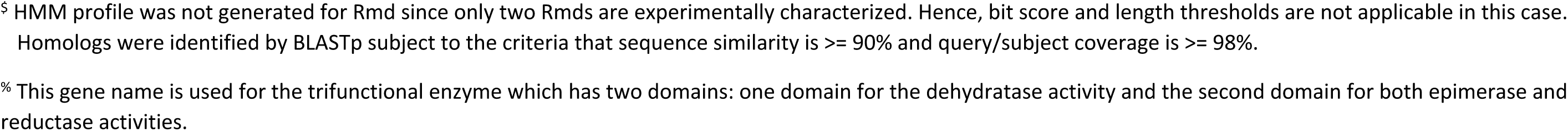
Keywords used for searching databases, number of sequences used for generating HMM profiles and thresholds†

| Profile | E. C. Number<sup>¶</sup> | Primary gene name<sup>¶</sup> | Primary enzyme activity (Annotation assigned to profile / BLASTp hits)<sup>¶</sup> | Number of proteins used for generating HMM profile |  |  |  | No. of proteins with known 3D structure | No. of NR sequences<sup>‡</sup> | Threshold<sup>#</sup> |  |
| --- | --- | --- | --- | --- | --- | --- | --- | --- | --- | --- | --- |
|  |  |  |  | Eubacteria | Archaea | Eukaryota | Virus |  |  | Bit score | Length (a.a.) |
| RmlA | 2.7.7.24 | *rmlA* | Glucose 1-phosphate thymidyltransferase | 42 | 3 | 0 | 0 | 12 | 31 | 162 | 219 |
| RmlB+Ugd | 4.2.1.46 | *rmlB* | (d)TDP glucose 4,6-dehydratase | 39 | 2 | 5 | 1 | 7 | 36 (32+4) | 223 | 199 |
|  | 4.2.1.76 | *RHM*<sup>%</sup> | UDP glucose 4,6-dehydratase | 0 | 0 | 6 | 1 | 0 |  |  |  |
| RmlC | 5.1.3.13 | *rmlC* | (d)TDP-4-dehydrorhamnose 3,5-epimerase<br>(d)TDP-4-dehydrorhamnose 3-epimerase | 24 | 2 | 0 | 0 | 12 | 19 | 140 | 131 |
| RmlD | 1.1.1.133 | *rmlD* | (d)TDP-4-dehydrorhamnose reductase | 19 | 2 | 0 | 0 | 5 | 16 | 189 | 204 |
| Uger | 5.1.3.- / 1.1.1.- | *RHM*<sup>%</sup> | UDP-4-keto-6-deoxy-glucose 3,5-epimerase / UDP-4-keto-L-rhamnose reductase | 0 | 0 | 8 | 1 | 1 | 4 | 193 | 271 |
| Gmd | 4.2.1.47 | *gmd* | GDP-mannose 4,6-dehydratase | 18 | 0 | 15 | 2 | 6 | 24 | 289 | 280 |
| Rmd<sup>§</sup> | 1.1.1.281 | *rmd* | GDP-4-keto-6-deoxy mannose reductase | 2 | 0 | 0 | 0 | 1 | 2 |  |  |

<sup>†</sup> Listing of all enzymes viz., UniProt IDs, PubMed ids, organism and strain names, gene names are given in the Profile dataset worksheet in the file rhamnose.xlsx.
<sup>¶</sup> These were used as keywords to search databases. Uger has two E.C. numbers since it is a bifunctional enzyme. For the enzymes of the rml pathway, no distinction is made between enzymes which use dTDP-sugar and TDP-sugar. For Uger, the gene name *UER* was also used. For Ugd and Uger, rhamnose synthase / synthetase were also used for enzyme activity.
<sup>‡</sup> This is the number of non-redundant sequences that were eventually used to generate HMM profile. 80% sequence identity cut-off was used as the redundancy criterion i.e., no two sequences within each dataset has more than 80% sequence identity. For the RmlB+Ugd profile, although numbers are shown separately for RmlB and Ugd, all sequences were used together to identify the non-redundant dataset.
<sup>#</sup> A database sequence is considered as a hit for the HMM profile only when it meets both the bit score and length thresholds. The latter is the minimum number of amino acid residues of the database sequence that should align with the HMM profile.
§ HMM profile was not generated for Rmd since only two Rmds are experimentally characterized. Hence, bit score and length thresholds are not applicable in this case. Homologs were identified by BLASTp subject to the criteria that sequence similarity is $\geq 90\%$ and query/subject coverage is $\geq 98\%$ .
% This gene name is used for the trifunctional enzyme which has two domains: one domain for the dehydratase activity and the second domain for both epimerase and reductase activities.

#### Assessment using the CATH database

CATH is derived from the PDB. Based on 3D structure similarities, protein domains are classified hierarchically as Class, Architecture, Topology, Homologous superfamily (Dawson et al., 2017). Members of a superfamily are separately classified in two distinct ways: structural clusters (based on structure similarity) and functional families (based on sequence similarity). All the sequences in the CATH database were scanned against the HMM profiles (CATH Validation sheet of rhamnose.xlsx). There were no false positives for any of the profiles as judged by CATH annotations (Table S2). Taking into consideration annotations of hits obtained from the Swiss-Prot, Pfam and CATH databases, it is seen that the profiles generated in this study have high specificity (Table S1-S2).

**Table 2.** Number of homologs of rhamnose biosynthesis pathway enzymes found in the TrEMBL database and genomes#

| Enzyme | Total number of hits | Current annotation of homologs in comparison to profile annotation* |  |  |  |  |  |  |  |  |
| --- | --- | --- | --- | --- | --- | --- | --- | --- | --- | --- |
|  |  | Annotated at molecular function level |  |  |  |  | Annotated only at biological process level | Annotated only at cellular component level | Existing annotation inadequate to reconcile | No annotation whatsoever |
|  |  | Matches | Fusion proteins with incomplete annotation <sup>¶</sup> | Does not match<sup>%</sup> | Consistent | Inconsistent |  |  |  |  |
| TrEMBL database |  |  |  |  |  |  |  |  |  |  |
| RmlA | 23965 | 21854 | 9 | 25+211 | 251 | 0 | 1310 | 0 | 16 | 289 |
| RmlB+Ugd | 22776 | 21890 | 7 | 29+3 | 114 | 65 | 9 | 0 | 3 | 656 |
| RmlC | 17881 | 17451 | 33 | 23+29 | 183 | 0 | 1 | 0 | 26 | 135 |
| RmlD | 14660 | 14570 | 3 | 1+0 | 23 | 0 | 0 | 0 | 4 | 59 |
| Uger | 941 | 167 | 19 | 92+4 | 103 | 5 | 0 | 0 | 4 | 547 |
| Gmd | 14302 | 13705 | 0 | 2+3 | 99 | 9 | 3 | 0 | 2 | 479 |
| Rmd | 20 | 9 | 0 | 4+0 | 0 | 6 | 0 | 0 | 0 | 1 |
| Genomes |  |  |  |  |  |  |  |  |  |  |
| RmlA | 4700 | 4673 | 0 | 7+0 | 7 | 0 | 0 | 0 | 0 | 13 |
| RmlB+Ugd | 6209 | 6070 | 0 | 2+1 | 39 | 5 | 2 | 0 | 0 | 90 |
| RmlC | 4392 | 3846 | 30 | 1+0 | 498 | 0 | 0 | 0 | 3 | 14 |
| RmlD | 4139 | 3860 | 11 | 0+3 | 248 | 0 | 0 | 0 | 5 | 12 |
| **Uger** | 752 | 479 | 0 | 174+0 | 32 | 1 | 0 | 0 | 0 | 66 |
| **Gmd** | 3275 | 3171 | 0 | 1+0 | 37 | 0 | 0 | 0 | 0 | 66 |
| **Rmd** | 18 | 8 | 0 | 1+0 | 1 | 8 | 0 | 0 | 0 | 0 |

### Identifying homologs of rhamnose biosynthesis genes from TrEMBL database

The database annotation matched with the profile annotation for more than 90% of the hits (except in case of Uger) thereby providing manual curation for these electronically annotated entries (Table 2). The remaining 10% of hits includes those with annotation at biological process level which is consistent with the profile annotation, no annotation whatsoever and incomplete molecular function annotation (consistent or inconsistent). These hits were (re)assigned profile annotation. Annotations of hits whose molecular function annotation does not match the profile annotation were resolved after error analysis, wherever possible, based on careful scrutiny. The details are given as supplementary material (Annotation resolution – TrEMBL and TrEMBL hits worksheets of rhamnose.xlsx). Only a small fraction of the ∼140 million entries of the TrEMBL database were found to be homologs of rml, udp and gdp pathway enzymes. The number of homologs of Uger is less compared to those of the rml pathway and Gmd. This can be attributed to the udp pathway being restricted to plants, algae and fungi, whose contribution to the TrEMBL database is just 10-14%.

Very few homologs of Rmd are present in the TrEMBL database in contrast to the number of homologs found for Gmd and rml pathway enzymes (Table 2). For reasons mentioned in Methods, criteria set for finding Rmd homologs are stringent. To determine if this stringency is the reason for finding fewer homologs, TrEMBL database was searched by relaxing the criteria i.e., a minimum of 70% query coverage and at least 50% sequence similarity: this resulted in additional 59 entries annotated as Rmd and having conserved GXXGXXG and YXXXK motifs. However, there were also a large number of false positives as judged by their annotations. Three of the Gmds characterized experimentally have Rmd activity also (15,41,42). Other Gmds haven’t been assayed for Rmd activity and hence, it is not clear whether all Gmds have rmd activity also or there are two classes of Gmds: with and without Rmd activity. Therefore, it may be envisaged that genomes in which Rmd homolog could not be found have bifunctional Gmds and thus, can carry out the function of Rmd too. It is also to be noted that the gdp pathway is for the biosynthesis of D-rhamnose and hence, finding fewer homologs can be suggestive of restricted usage of D-rhamnose.

### Homologs of rhamnose biosynthesis genes from genomes

The number of hits obtained against genomes are tabulated in Table 2. Comparison and reconciliation of annotations were carried out as done for TrEMBL hits (Details given in Annotation resolution – Genomes and Genome hits worksheets of rhamnose.xlsx). As in the case of TrEMBL hits, annotations of most of the hits match corresponding profile annotation. Ugd and Uger are fused in 111 genomes. The presence of functionally interacting proteins as fused proteins is not unexpected and Ugd and Uger activities are known to be present in a single polypeptide chain in many plants, e.g., *Arabidopsis thaliana* (Oka, Nemoto, & Jigami, 2007). In contrast, only around 1% of RmlC and RmlD hits occur as fused proteins; in case of RmlA and RmlB hits, only 1 or 2 instances of fused proteins were found. In *Desulfurivibrio alkaliphilus* AHT 2 and *Rhodococcus* sp. P1Y, RmlA and RmlC hits occur as fused proteins. Even though, RmlA and RmlC are not consecutive in the pathway, these genes occurring as a fusion protein is not unusual: the bifunctional *E. coli* protein HldE catalyses non-consecutive steps in the biosynthesis of ADP-L-*glycero*-β-D-*manno*-heptose (43). Two sets of rml genes are present in *Desulfurivibrio alkaliphilus* AHT 2. The order of the two sets is different: (i) RmlC, RmlA, RmlD and RmlB at one locus and (ii) fused RmlA-RmlC, RmlD and RmlB at the other. Several species of *Rhodococcus* have been sequenced, but RmlA and RmlC are fused in only one.

### Presence of rhamnose biosynthesis pathway in genomes

Presence of homologs for all enzymes of the rml, gdp or udp pathway suggests that the organism uses rhamnose as a glycan building block. Overall, it is found that the rml pathway, the most prevalent of the three, is found only in 42% and 21% of bacterial and archaeal genomes, respectively (Table 3). The udp pathway is found in 18% of eukaryotic genomes. The gdp pathway is found mostly in *Pseudomonas*. Gmds from some organisms e.g., *Paramecium bursaria Chlorella* virus 1 are bifunctional i.e., they have Rmd activity also (42). The gdp pathway will be functional in many other genomes wherein only Gmd homolog is found but not that of Rmd, if such Gmd homologs are also bifunctional.

**Table 3.** Prevalence of rhamnose biosynthesis pathways

| Domain | Total number of genomes | Number of genomes |  |  |  |  |  |  |  |  |
| --- | --- | --- | --- | --- | --- | --- | --- | --- | --- | --- |
|  |  | rml pathway |  |  | gdp pathway |  |  | udp pathway |  |  |
|  |  | Complete pathway present | Partial pathway present | Pathway absent | Complete pathway present* | Partial pathway present | Pathway absent | Complete pathway present | Partial pathway present | Pathway absent |
| Bacteria | 12823 | 5377 | 4500 | 2946 | 162 | 4452 | 8209 | 0 | 9120 | 3703 |
| Archaea | 303 | 65 | 115 | 123 | 0 | 39 | 264 | 0 | 148 | 155 |
| Eukaryota | 993 | 0 | 508 | 485 | 0 | 579 | 414 | 186 | 329 | 478 |
| Virus | 7845 | 2 | 8 | 7835 | 0 | 19 | 7826 | 1 | 9 | 7835 |
| Total | 21964 | 5444 | 5131 | 11389 | 162 | 5089 | 16713 | 187 | 9606 | 12171 |

\* 161 out of 162 genomes also have the rml pathway and all these belong to the genus *Pseudomonas*. *Aneurinibacillus* sp. XH2 genome does not have the rml pathway.

All genes of a pathway are contiguous in the genome only in some organisms (Table 4), suggesting that the expression of genes is co-regulated in such genomes. The rml pathway genes are contiguous in all strains of *Pseudomonas aeruginosa*, *Burkholderia pseudomallei*, *Enterococcus faecalis*, *Sinorhizobium meliloti*, *Porphyromonas gingivalis*, etc., whereas they are dispersed in all strains of some organisms like *Neisseria gonorrhoeae*, *Streptococcus thermophilus*, *Mycobacterium avium*, etc.; they are in the neighbourhood of each other in all strains of organisms like *Lactococcus lactis*, *Phaeobacter inhibens*, etc. The genes of the rml pathway are found to be dispersed in *Mycobacterium tuberculosis* H37Rv (37). In fact, these genes are dispersed in 172 of the 174 strains of *M. tuberculosis* analysed in this study. RmlD homolog is missing in the remaining two strains, namely Haarlem/NITR202 and CAS/NITR204.

**Table 4.**
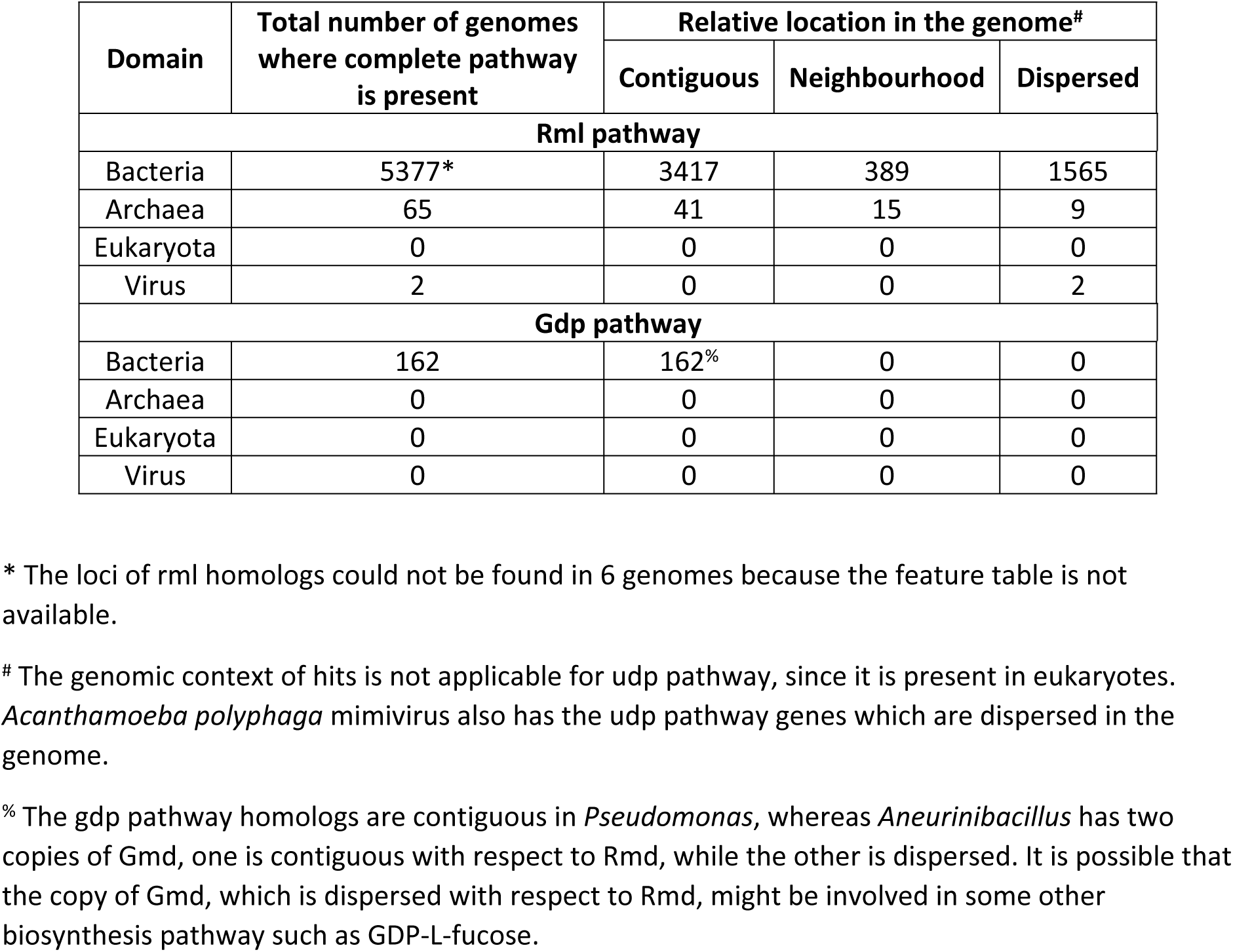
Relative location of the rhamnose biosynthesis genes in the genome

One or more genes of the biosynthesis pathway are missing in some of the genomes (Table 3). It is possible that RmlA and RmlB homologs are a part of some other biosynthesis pathway e.g., dTDP-fucose in genomes wherein RmlC and RmlD homologs are missing. Similarly, RmlA, RmlB and RmlC homologs are involved in the biosynthesis of sugars such as dTDP-6-deoxy-L-talose in genomes wherein RmlD is missing. One can envisage the presence of a broad specificity nucleotidyltransferase in genomes which have RmlB, RmlC and RmlD homologs but not that of RmlA e.g., *Bacillus anthracis* str. Ames. There are several genomes in which only Gmd homolog is found; it is possible that such Gmd homologs do not have Rmd activity but are a part of a pathway for the biosynthesis of other sugars such as GDP-L-fucose. It is also possible that homologs, even when present, are not identified by HMM profiles because of sequence divergence / threshold stringency.

Furthermore, in some genomes, more than one hit for rhamnose biosynthesis genes is found (Genomes worksheet of rhamnose.xlsx). It is possible that the other copy of these genes might be involved in some other pathway, eg., TDP-Fuc4NAc biosynthesis pathway (44) or might be a result of gene duplication event. In most of the organisms, rhamnose biosynthesis genes are present on the chromosome; in some organisms which have two sets of these genes, one set is present on the chromosome and the other set on the plasmid, e.g., as in *Bacillus thuringiensis* serovar *indiana* strain=HD521. Similarly, some organisms have only one set of these genes which is distributed on the chromosome and plasmid, eg., *Paludisphaera borealis* strain=PX4 which has RmlA and RmlC on the plasmid, and RmlB and RmlD on the chromosome.

### Prevalence of the rhamnose biosynthesis pathway at various levels of classification

The rml pathway is found in bacteria, archaea and phages but not in eukaryotes or other viruses. The udp pathway is found only in plants, fungi, algae, nematodes, a chordate and *Acanthamoeba polyphaga* mimivirus. The gdp pathway is found only in *Aneurinibacillus*, *Pseudomonas* and a virus (chlorovirus PBCV-1). Organisms which have the udp pathway do not have the rml or gdp pathway and vice versa. 154 of the 175 strains of *Pseudomonas aeruginosa*, *Pseudomonas fluorescens* (strain=NCTC10783) and *Pseudomonas* sp. AK6U have both gdp and rml pathways (Table 3 and Genomes sheet of rhamnose.xlsx). Rml pathway genes are not found in *Aneurinibacillus* sp. XH2, which is seen to have the gdp pathway.

The prevalence of the three pathways within different phyla (with at least 50 sequenced genomes) varies significantly. Actinobacteria, Bacteroidetes, Crenarchaeota, Cyanobacteria, Fusobacteria and Proteobacteria are phyla in which rml pathway is predominant (at least 40% genomes). In contrast, <20% organisms of Chlamydiae, Euryarchaeota and Tenericutes have the rml pathway. The udp pathway is found in all genomes of Streptophyta, <=25% genomes of Ascomycota and Chordata, and none of the genomes of Arthropoda, Basidiomycota, Chlamydiae and Negarnaviricota.

The presence of rhamnose has not been reported in *Chlamydia, Helicobacter, Mycoplasma, Pasteurella, Rickettsia* and *Staphylococcus*, although all these have lipopolysaccharides and cell-surface appendages. Expectedly, none of the rhamnose pathways are found in this study. All these are pathogenic including *Pasteurella*, a zoonotic pathogen transmitted to humans by domestic animals. *Mycoplasma* is devoid of cell-wall but, nevertheless, is known to synthesise lipopolysaccharides. *Rickettsia* and *Pasteurella* are non-motile, pleomorphic organisms.

There are in total, 3,960 species included in the dataset. Only 29 of these 3,960 species have >50 strains (Figure S3). Strain-specific variations with respect to the presence of rml pathway are mainly found in pathogenic strains of organisms such as *Escherichia coli*, *Salmonella enterica*, *Klebsiella pneumoniae*, *Mycobacterium tuberculosis*, *Streptococcus pyogenes*, *Acinetobacter baumannii*, *Neisseria meningitidis*, *Streptococcus pneumoniae*, *Xanthomonas citri* (a phytopathogen) and *Clostridioides difficile* (Prevalence of pathways worksheet of rhamnose.xlsx).

The rhamnose biosynthesis pathway is notably absent in chordate genomes which includes mammalian (144), avian (72) and ichthian (64) genomes. The exception is Chinese rufous horseshoe bat which has three homologs each of Ugd and Uger. Of these three, one is a Ugd-Uger fusion protein (XP_019575949.1). The bit scores of these homologs are much higher than threshold implying that they are unlikely to be false positives. Moreover, functionally important residues are conserved in these homologs. Rhamnose biosynthesis pathways are not found in any of the other six bat genomes included in the dataset. However, the role of rhamnose in this bat species is not clear.

### Genomic context of rhamnose biosynthesis pathway genes

As mentioned earlier, rml and gdp pathway genes are contiguous or in neighbourhood of each other in some of the genomes. In such genomes, functional annotations of flanking genes were analysed with a view to determine the biological processes in which rhamnose might be used. In some genomes, rml pathway genes are flanked by genes involved in CPS/EPS/LPS/Capsid biosynthesis, secondary metabolite production, and cell surface appendage biosynthesis and modification (Figure 2). In a few other genomes, the biological functions of flanking genes include other related functions such as the biosynthesis of quinovosamine, a glycan building block found in LPS and flagellin. Even in the case of gdp pathway genes, biological functions of flanking genes are related (Figure 2). The rml / gdp pathway genes in these genomes can be additionally annotated at the biological process level also. The three biological processes are not mutually exclusive, meaning an organism can have both CPS/EPS/LPS and cell surface appendages and synthesize secondary metabolites. Therefore, the synthesized rhamnose can be utilised in all or most of these biological processes in an organism. However, the regulation of expression of rhamnose biosynthesis genes in these cases might be independent from the regulation and expression of genes related to the biological processes mentioned above.

**Fig. 2.**
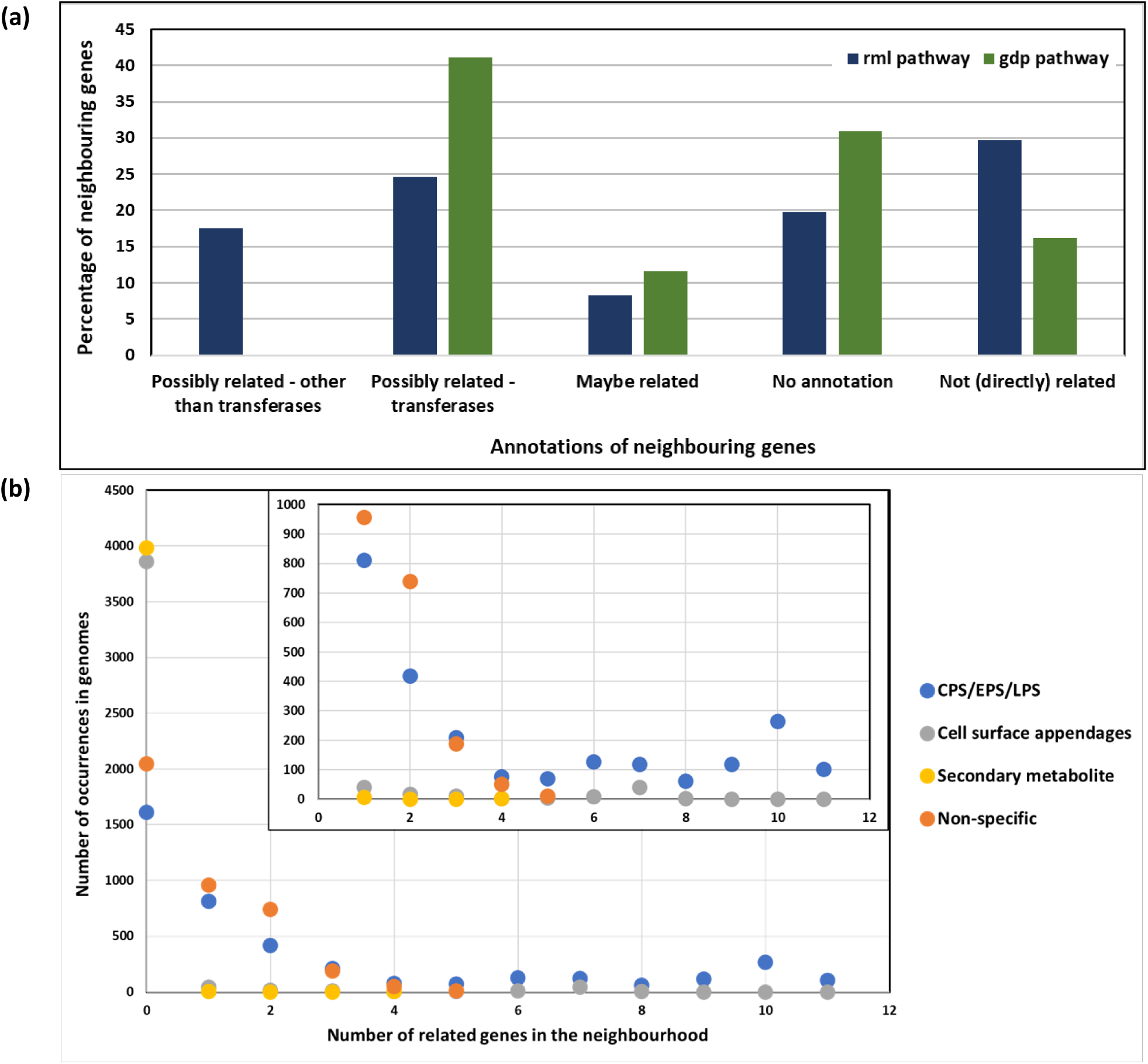
Genomic context of rml and gdp pathway genes as determined by annotations of neighbouring genes. Genes are categorized on the basis of the (probable) biological function that the gene might be involved in. The genomic context of the udp pathway was not analysed since this pathway is found only in eukaryotes. (a) The “Possibly related – other than transferases” category includes genes whose annotations are related to the following: (i) CPS/EPS/LPS/Capsid biosynthesis, (ii) secondary metabolite production and (iii) cell surface appendage biosynthesis and modification. The “Possibly related – transferases” category includes glycosyl- and methyl-transferases. The “Maybe related” category includes genes annotated as epimerases, dehydratases, hydratases, etc without any specification of the substrate. (b) Sub-grouping of genes in the “Possibly related – other than transferases” category. The “non-specific” category includes genes which might be involved in multiple biological functions such as genes involved in the biosynthesis of 4-aminoquinovosamine which are involved in both LPS biosynthesis and flagellin modification. [Inset] Zoomed in view.

### Prevalence of rhamnose biosynthesis pathways in some common genera and species

#### Pseudomonas

Lipopolysaccharides (LPS) in *Pseudomonas* are structurally diverse (Lam, Taylor, Islam, Hao, & Kocíncová, 2011; McCaughey et al., 2014; Ovod, Rudolph, Knirel, & Krohn, 1996). The two major types are CPA (common polysaccharide antigen) and OSA (O-specific antigen). CPA is a homopolymer of D-Rha with the repeating unit [-D-Rha-α1,2-D-Rha-α1,3-D-Rha-α1,3-]. OSA is devoid of rhamnose. CPA and OSA are linked to the core structure through an L-Rha residue. Thus, both enantiomers of rhamnose are utilised in *Pseudomonas* (47). How CPA and OSA are linked to the core oligosaccharide in organisms which lack rml pathway and what substitutes for CPA in organisms which lack the gdp pathway are not clear. Mutation studies in *P. aeruginosa* PAO1 have shown that inability to synthesize D- and L-Rha results in the formation of incomplete core which lacks both CPA and OSA (31). Some of the species which show strain-specific presence / absence of rhamnose biosynthesis pathways are *P. aeruginosa, P. chlororaphis, P. putida* and *P. fulva*. Knowledge of phenotypic characteristics and ecological niche of these organisms will help in understanding the role of rhamnose.

#### Acinetobacter

Rhamnose is a constituent of capsular O-antigen of *A. baylyi* (also known as *A. calcoaceticus*) BD4 and BD413 (ADP1) (48–51). Expectedly, the rml pathway is found in 34 strains of *Acinetobacter* including *A. baylyi* ADP1 (*Acinetobacter* sp. ADP1); genome sequence of *A. baylyi* BD4 is not available. The rhamnose-glucose capsule of BD4 has been reported to interfere with adherence to hydrocarbons and thus, decrease cell surface hydrophobicity (51). Among *Acinetobacter baumannii* strains, rhamnose is found in serogroup O10 (52), serogroup O-7 (53) and NCTC 10303 (54), but not in strains 24, 34, 108 and 214 (55–57). However, genome sequences are not available for any of these strains. Rml pathway is found in only 12 of the 128 strains of *Acinetobacter baumannii* (Figure 3) for which genome sequences are available. Experimental data for the absence of rml pathway is available only for the strain 307-0294 (58).

**Fig. 3.**
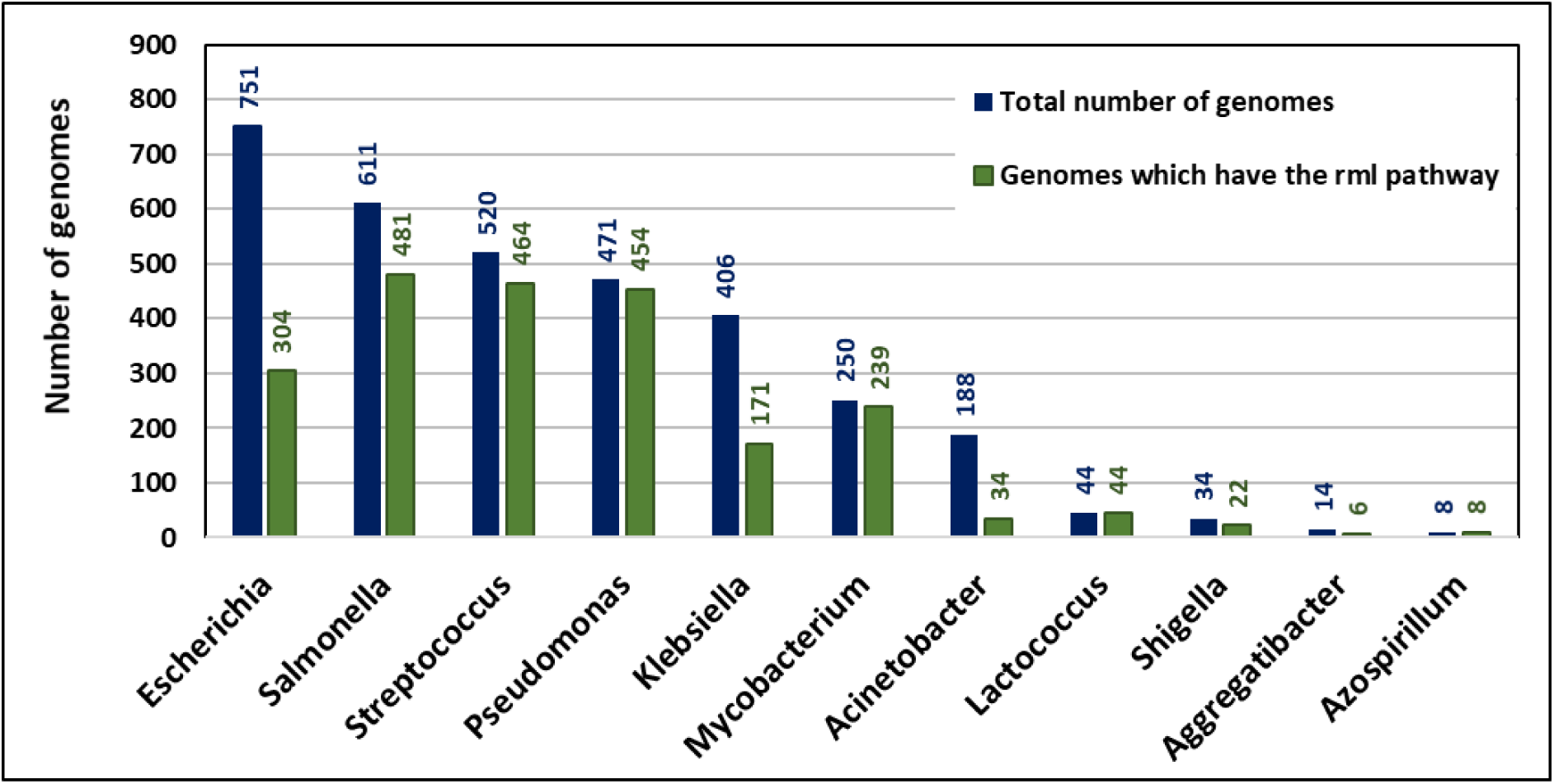
Genus-specific variation in the prevalence of the rml pathway.

#### Azospirillum brasilense

Rhamnose is a part of LPS and disruption of rhamnose biosynthesis genes results in modified LPS core structure, non-mucoid colony morphology, increased EPS production, and also affects maize root colonization (59). Rml pathway genes are found in both strains for which genome sequence is available.

#### Aggregatibacter actinomycetemcomitans

The presence of rhamnose is strain-specific in this species (60, 61). Rml pathway is found in 6 of the 8 strains for which genome sequence is available (Figure 3). It is absent in the strains D7S-1 and 624 in agreement with the literature reports (62) (63).

#### Klebsiella

The O12 antigen of K. pneumoniae is said to be composed of an N-acetylglucosamine and rhamnose polymer (64). The rml pathway is found in 133 of the 320 strains of *Klebsiella pneumoniae*. However, genome IDs could not be mapped to serotypes in which the rhamnose pathway genes are known to be a part of the cps cluster (65–71). The sole exception is *K. pneumoniae subsp. pneumoniae* MGH 78578 which is mapped to serotype K52 and in this strain rml genes are contiguous (Genomes worksheet of rhamnose.xlsx).

#### Lactococcus and Streptococcus

The presence of rml pathway is strain-specific in *S. pyogenes, S. mutans, S. suis* and *S. pneumoniae*, but is found in all strains of *L. lactis* and *S. thermophilus* (Genomes worksheet of rhamnose.xlsx). dTDP-L-rhamnose is an important precursor of CPS and EPS in lactococci, lactobacilli and streptococci (1, 72). The rml pathway genes have been found to be essential in *L. lactis* MG1363 (6). Rml genes have been characterized and appear to play a vital role in the production of serotype-specific, rhamnose-containing CPS antigens in *S. mutans* (73, 74) and *S. pneumoniae* (75–77). Rml mutations in *S. mutans* resulted in a change in the composition of CPS and absence of the serotype-specific O antigen. *S. pneumoniae* cps19fL-cps19fO (RmlA-RmlD) mutants exhibited a so-called rough non-encapsulated phenotype and did not have the capacity to produce CPS, indicating that the rfb analogues play an essential role in CPS-19F production (78).

#### Mycobacterium

L-rhamnose covalently links arabinogalactan to peptidoglycan (79), which is critical for the overall architecture of the mycobacterial cell wall, making L-rhamnose biosynthesis essential (36, 80). Rml genes are dispersed in mycobacterial genome (37). L-rhamnose biosynthesis pathway is considered a promising drug-target, since this pathway is absent in humans (35, 81). The rml pathway is found in all 22 strains of *M. avium* and in 172 strains (out of 174) of *M. tuberculosis.* The homolog for RmlD is missing in other 2 strains (strain Haarlem/NITR202 and CAS/NITR204). Only one report for these strains is available (82) and otherwise not much information is available for these strains.

#### Shigella

The rml pathway is fairly conserved in *Shigella* species and is a common constituent of the O-antigen structure (10,83,84). Several rhamnosyltransferases have also been reported for *Shigella dysenteriae*. The rml pathway is observed in 15 strains (out of 16) of *S. flexneri* and 3 strains (out of 6) of *S. dysenteriae* (Genomes worksheet of rhamnose.xlsx).

#### Escherichia

Among the 751 *Escherichia* genomes, 304 genomes have all the rml pathway enzymes (Figure 3). None of these genomes have gdp or udp pathway. *E. fergusonii* ATCC 35469 and *E. marmotae* HT073016 have partial rml pathway. Of the 14 strains of *E. albertii*, only three have the complete rml pathway (strains: KF1, NIAH_Bird_3 and 2012EL-1823B); others have partial rml pathway. 301 strains, out of 735 strains of *E. coli*, have the complete rml pathway and 32 strains do not have any rml pathway enzymes. L-rhamnose has been reported to a part of K- and O-antigens in *Escherichia* (11,85–88). Burns and Hull in 1998 demonstrated the importance of rhamnose biosynthesis pathway in *E. coli.* Mutation of RmlD resulted in loss of O-antigen expression but leaves the lipopolysaccharide core and lipid A structure intact, yielding viable bacteria. However, rhamnose-deficient *E. coli* are extremely sensitized to serum mediated killing (34).

#### Salmonella

Out of the total 611 genomes included in the dataset belonging to *Salmonella* genus, 605 are strains of *S. enterica*. 479 strains of *S. enterica* have the complete rml pathway; the remaining 126 strains have partial rml pathway. The O-antigen of *S. enterica* serovar Typhimurium consists of a repeat unit of four sugars: abequose, mannose, rhamnose and galactose (9). The O-antigens form hydrophilic surface layers that protect the bacterium from complement-mediated cell lysis (89). All the four strains of *S. bongori* (a species that is predominantly associated with cold-blooded animals, but in rare cases, infects humans also) have partial rml pathway, implying the rml genes might be involved in some other biosynthesis pathway, other than rhamnose biosynthesis. *Salmonella* sp. SSDFZ54 and SSDFZ69 have the rml pathway, however no literature is available for these strains.

### Homologs of rhamnose biosynthesis genes in human-associated and environmental metagenomes

Analysis of human-associated metagenomes showed the presence of all 4 genes of the rml pathway in a large number of samples. Analysis of environmental metagenomes showed that the rml pathway genes are found in all samples from grasslands, lake, sand and sediment, and in >30% of samples from glacier, hydrothermal vents, marine, oceanic, photic zone and soil. The udp pathway is found only in 6 samples of human faecal metagenome (PRJNA354235) (Metagenome hits and Human-associated metagenome hits worksheets of rhamnose.xlsx). The presence of organisms belonging to Streptophyta and Ascomycota in these samples has been reported (https://www.ebi.ac.uk/metagenomics/studies/MGYS00003733#analysis). Therefore, the presence of plant, algae and / or fungal genomic content in the faecal samples can be inferred based on the presence of the udp pathway in these samples. The udp pathway is found in very few environmental metagenome samples; these samples are from coastal, marine, salt marsh and sand biomes (Metagenome hits and Environmental metagenome hits worksheets of rhamnose.xlsx). No homologs for Rmd were found in human-associated and environmental metagenomes. This maybe because of (i) stringency of the parameters set for identification of Rmd homologs, or (ii) absence of bacteria synthesising GDP-D-rhamnose in the samples included in the dataset. However, it is possible that the Gmd homologs found in the human-associated and environmental metagenomes might be bifunctional.

The composition of organisms at the level of phylum, class, family, genus or species has been reported for some of the human-associated and environmental biomes. The presence of the rml pathway genes in genomes belonging to these phyla, class, etc. (Genomes worksheet of rhamnose.xlsx) were checked. It is found that the two data are in consonance with each other. For e.g., in the gut and oral microbiome of individuals with rheumatoid arthritis, *Haemophilus spp*. are depleted and *Lactobacillus salivarius* is over-represented; the genome data suggests that only one strain of *Haemophilus* and 6 out of 8 strains of *Lactobacillus salivarius* have the rml pathway which might be contributing to the presence of rhamnose biosynthesis genes in the biome Table S3.

## DISCUSSION

Glycosylation is found in all domains of life, i.e., eukaryotes, archaea, eubacteria and viruses. S-layer glycoproteins, pili, flagella, fimbriae and secreted glycoproteins are some examples of prokaryotic glycoconjugates which play an important role in cell physiology, microbe–host interactions, immune escape mechanisms and biofilm formation. Prokaryotic glycans seem to show far more structural diversity than eukaryotic glycans; even serotype-specific differences are known. However, not much is known, especially in prokaryotes, about molecular details of glycan biosynthesis and factors that govern structural diversity. Consequently, establishing the relationship between structure and function has become very hard. This is in stark contrast to proteins where relating sequence changes to functional changes via site-directed mutagenesis has become routine.

Rhamnose is a typical constituent of capsular polysaccharides and glycan moieties of flagella, fimbriae and pili. It is also a precursor for secondary metabolites such as streptomycin in bacteria (90) and flavonoids in plants (3, 5). So far, rhamnose has not been found in any chordate. Pathways for the biosynthesis of rhamnose have been elucidated in a few organisms. This knowledge is exploited in the present study to investigate the prevalence of rhamnose biosynthesis pathways in organisms with completely sequenced genomes.

Only a small fraction of protein sequences deposited in the databases have experimentally proven biological activity. The molecular function of a protein is often inferred on the basis of its sequence similarity to a protein whose function has been experimentally determined. In this study, protein family-specific profiles for rhamnose biosynthesis enzymes were generated by including such experimentally characterised proteins and the specificity and sensitivity of these profiles was put to test by running them against curated sets of proteins from Swiss-Prot, Pfam and CATH databases. HMM profile for Rmd could not be generated due to lack of many experimentally demonstrated Rmds. In this case, a BLASTp approach was taken to identify homologs of Rmd.

The presence of rhamnose, though demonstrated to be important in virulence of several bacteria (30,32,33), is not limited to pathogenic bacteria. There are many non-pathogenic bacteria which also have rhamnose such as *Lactobacillus* and *Lactococcus*. Also, rhamnose is found in intracellular (e.g., *Lawsonia intracellularis*), facultative intracellular (e.g., *Mycobacterium tuberculosis*) and extracellular organisms (e.g., *Streptococcus pyogenes*). Rhamnose is found in both aerobic (e.g., *Mycobacterium tuberculosis*) and anaerobic bacteria (e.g., *Prevotella*).

The rml pathway is observed in bacteria, archaea and phages but not in any of the eukaryotes. *Burkholderia*, *<u>Enterococcus</u>*, *<u>Lactococcus</u>*, *Mycobacterium*, *Salmonella*, *Streptococcus*, *Sulfolobus* and *<u>Xanthomonas</u>* are the most prominent genera which have the rml pathway. This pathway is relatively less prevalent in *<u>Acinetobacter</u>*, *Bacillus*, *<u>Bordetella</u>*, *<u>Clostridium</u>*, *<u>Corynebacterium</u>*, *Escherichia*, *<u>Lactobacillus</u>*, *Neisseria*, *Streptomyces* and *<u>Vibrio</u>*. *Bordetella* are pathogenic obligate aerobes, lactobacilli live in human digestive, urinary and genital systems, *Corynebacterium* are commensal aerobes and Escherichia may be free-living or found in GI tract. Genera underlined as those in which the rml pathway has not been characterized so far. Key genera where this pathway is absent include *Bifidobacterium*, *Chlamydia*, *Helicobacter*, *Listeria*, *Mycoplasma*, *Pasteurella*, *Rickettsia* and *Staphylococcus*; many of these are obligate or facultative intracellular organisms and the relationship between this niche and absence of rhamnose needs further exploration. The gdp pathway, on the other hand, is only found in *Pseudomonas* and *Aneurinibacillus*. It is not clear why Pseudomonas uses both D- and L-enantiomers of rhamnose (biosynthesized through the gdp and rml pathways, respectively), instead of using only one enantiomer like all other organisms, is not clear.

At the level of species, the rml pathway is highly prevalent in *<u>Enterococcus faecalis</u>*, *<u>Enterococcus faecium</u>*, *Mycobacterium avium*, *Mycobacterium tuberculosis*, *Neisseria gonorrhoeae*, *Pseudomonas aeruginosa*, *Pseudomonas syringae*, *Salmonella enterica*, *Shigella flexneri*, *Streptococcus pyogenes*, *Streptococcus suis* and *Streptococcus thermophilus*. The rml pathway is absent in *Bordetella pertussis*, *Campylobacter jejuni*, *Bacillus subtilis*, *Bacillus anthracis* and *Vibrio cholerae*.

The udp pathway is found only in plants, algae, fungi, nematodes, a chordate (Chinese rufous horseshoe bat) and a virus. The presence of the udp pathway only in this bat genome but none of the other 302 chordate genomes is baffling. Flavonoids are widely distributed in plants and fulfil several functions (91). L-rhamnose is a part of pectin, a heteropolysaccharide present in the primary cell walls of terrestrial plants (92). According to the present study, rhamnose is observed to be present in all plants (Streptophyta) and most of them are known to have pectin cell wall and produce flavonoids. It has been suggested Ugd and Uger have been acquired by viruses through horizontal gene transfer and that these enzymes are involved in posttranslational modification of capsid proteins (93).

Some of the approaches to assign biological process level annotation rely upon genomic context since co-occurrence of genes suggest functional interaction. However, these approaches are ineffective in cases such as the rhamnose biosynthesis genes. This is because rhamnose is used by organisms for multiple functions e.g., CPS/EPS/LPS and cell surface appendages and synthesize secondary metabolites.

Large volumes of genomic and transcriptomic data have been generated but deciphering the biological information hidden in these data is a challenge. A significant portion of identified proteins remain annotated as proteins or domains with unknown function (PUFs and DUFs). Typical enzyme activity and binding assays are low-throughput “one protein at a time” approaches but they provide high quality molecular function level annotation. Experimental techniques are constrained by time and cost, and are not practically possible in all cases. To overcome this, various electronic annotation tools which annotate a protein at the molecular function level have been developed. In a large number of cases, proteins which share (high) sequence similarity perform the same molecular function. But electronic annotations are “putative” until proven experimentally. Such annotations may be incomplete or fully/partly erroneous. It is also known that some proteins with similar sequences catalyse the same reaction but with different substrates. Additionally, molecular function of a protein can be assigned with varying levels of details. In general, the extent to which sequence similarity translates to similarity in molecular function varies and this poses a challenge for electronic annotation.

Metagenomics has turned out to be an important tool in unravelling microbial communities of biomes. Understanding the mutual influence of microbiota and metabolism in humans has been the focus of several studies. Towards this, identification of operational taxonomic units (OTUs) that constitute a biome is important. Determination of surface polysaccharide determinants and biosynthesis pathways can facilitate relatively more fine-grained identification of OTUs. Rhamnose biosynthesis pathway can be one such determinant because it is a common constituent of surface polysaccharides and glycoproteins.

## MATERIALS AND METHODS

### Databases and software tools

Protein sequences are from UniProtKB, PDB, NCBI, Pfam and CATH databases (Table S4). Software tools available in the public domain (Table S4) were installed and executed locally. In addition, Python, Perl, R and shell scripts were written in-house as per needs. Default values were used for parameters of BLASTp and HMMER, unless stated otherwise.

Sequences of the full complement of proteins encoded by genomes are from the NCBI database (March 2019 release). This included 12,823 bacterial, 303 archaeal, 7,845 viral and 993 eukaryotic (includes contigs and scaffolds) genomes (the number of viral genomes that encode >1000 and >500 proteins is 5 and 20, respectively). The total number of unique proteins from these 21,964 genomes is 45,132,458. Feature tables required to analyse the genomic context of homologs from eubacteria, virus and archaea are from NCBI. GenBank equivalents were used for 17 bacterial genomes for which feature tables are not available. Even GenBank feature table was not available for *Peptoclostridium difficile* BI1 (GCF_000211235.1) genome. Protein sequences from WGS metagenome assemblies, and their BioSample, BioProject and Biome information are from the MGnify resource of EBI (Table S4). The corresponding BioProject metadata were extracted from the SRA section of NCBI. Gene ontology consortium’s prescription of assigning functions to a protein at three levels, namely, molecular function, cellular component and biological process, has been adapted in this study.

### Procedure used for generating HMM profiles

All proteins that catalyse the same step of a pathway (Figure 1) were considered to form a functional family. For each functional family, a family-specific HMM profile was generated as follows: PubMed, SwissProt and PDB were searched using EC number and primary gene name as keywords (Table 1). Corresponding original research articles were read to prepare a carefully curated set of experimentally characterized proteins. All-against-all pairwise alignments were used to ensure that proteins are sequence homologs. Query and subject coverages were used to identify relevant domains in case of multi-domain or fusion proteins. A sequence identity cut-off of 80% was applied to obtain a non-redundant dataset and this was designated as the Exp dataset; here, Exp signifies that the molecular function of all entries of this dataset has been experimentally characterized by direct activity assay. A multiple sequence alignment of Exp dataset proteins was used to generate a HMM profile, designated as the Exp profile. Exp dataset proteins were scored against the Exp profile and the floor value of bit score of the lowest scoring protein was set as the threshold for the Exp profile. “Best 1 domain” bit scores (T_exp bits) were used instead of E-values to facilitate comparison independent of the size of the search space.

With a view to enhance HMM profile’s coverage of sequence divergence within a functional family, an extended dataset (referred to henceforth as Extend dataset) was created starting from the Exp dataset by adding proteins that meet any of the following criteria:

i. Swiss-Prot entries that score >= T_exp against the Exp profile.
ii. Swiss-Prot entries that score < T_exp but whose annotation is consistent with the molecular function of the corresponding functional family.
iii. PDB entries whose annotation is consistent with the molecular function of the functional family but direct activity assay has not been reported. The protein annotated as “NafoA.00085.b” (A0A1W2VMZ8) was included in the Gmd profile since it has residue conservation.
iv. Proteins which have been inferred to have the molecular function by complementation assays or product characterization.

In case of (ii), (iii) and (iv), binding and catalytic site residues were ascertained to be conserved. Functionally important residues were identified as follows:

a. From the literature e.g., data based on site directed mutagenesis.
b. Multiple sequence alignment of Exp dataset proteins.
c. From 3D structures i.e., those that form H-bond with the ligand through side chain (Table S5).

A multiple sequence alignment of Extend dataset proteins was used to generate the Extend HMM profile. Details of proteins viz., source organism, UniProt ID, PubMed ID (for proteins for which experimental data is available) and gene name are given in the worksheet named as Profile Dataset (electronic supplement file rhamnose.xlsx) and a summary is given in (Table 1).

### Setting bit score thresholds for Extend profiles

Several of the entries in the TrEMBL database have UniRule / SAAS based electronic annotations (94). UniRule uses Pfam, TIGRFam and ProSite signatures along with some additional constraints to scan and annotate sequences. These signatures comprise of conserved residues of a protein family. These annotations were made use of to generate ROC curves, based on which bit score thresholds were set for Extend profiles (Figure S4). TrEMBL database was searched by varying the bit score threshold. For each bit score threshold, from among the hits, proteins which have molecular function annotation were sorted as true positives (TP) or false positives (FP) depending upon whether the annotation is consistent (TP) or inconsistent (FP) with the profile annotation. The profile annotation is the representative annotation of the protein family. From among the non-hits, false negatives (FN) were identified if the molecular function annotation of the entry is consistent with that of the Extend dataset proteins. True positive rate and false positive rate were calculated as 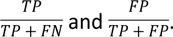

### Setting length threshold

Some of the entries in databases are not full-length proteins but are either fragments or truncated proteins. To filter out such entries, a length threshold also was set for each Extend profile. This is the shortest length of the multiple sequence alignment of Extend dataset proteins that covers all conserved regions. The length threshold is applied to the alignment length of the protein with the HMM profile, rather than the full-length of the protein. Database entries satisfying both bit score and length thresholds were analysed by multiple sequence alignment for conservation of functionally important residues. In this step, residues forming H-bonds through main chain were ignored and conservative replacements were allowed except for catalytic site residues. Proteins satisfying all three criteria viz., bit score and length thresholds of Extend profiles (Table 1) and conservation of key residues (Table S5) were considered as hits for that HMM profile i.e., functional homologs.

Applying the residue conservation criterion resulted in the elimination of 7-30% of the hits obtained from HMM profiles (Table S6). It is possible that some of these hits have intragenic compensatory mutations that will restore activity. It is also possible that the enzyme activity is lost in these homologs. Applying the length criterion further increased the specificity of search for homologs primarily in the case of Uger and Gmd; these homologs are fragments, not full-length sequences.

### Functional family-specific details pertaining to HMM profile generation

Ugd and Uger form a fusion protein, especially in plants. These two domains were taken separately for generating HMM profiles. Both dTDP-L-rhamnose and TDP-L-rhamnose are used as donor substrates during glycan biosynthesis. Literature reports suggest that enzymes of the rml pathway can utilize both ribo- and deoxyribo-substrates. The extent of specificity may vary for enzymes from different organisms. In fact, dTDP and TDP have been used synonymously by some authors. In view of this, no effort is made in the present study to distinguish these two nucleotide sugars.

#### RmlA

*Sulfolobus tokodaii* RmlA (Q975F9) is unique among RmlA in having an LβH domain at the C-terminus; however, the entire sequence was used for generating the HMM profile.

#### RmlB and Ugd

Both these are glucose 4,6-dehydratases except that RmlB uses (d)TDP-glucose as the substrate whereas Ugd uses UDP-glucose. Experimentally characterized RmlBs and Ugds (only the dehydratase domain, in case of multi-domain Ugd) align end-to-end and share 50-67% sequence similarity. HMM profiles generated using only RmlBs or only Ugds were not able to discriminate these two enzyme classes. Hence, a single RmlB-Ugd profile was generated; hits will be treated as (d)TDP/UDP-glucose 4,6-dehydratases.

#### RmlC

The gene name *rmlC* is associated in literature with two very similar, but distinct, enzyme activities: 3,5-epimerisation or 3-epimerisation of (d)TDP-4-keto-6-deoxy glucose (Figure S5). Of the 12 RmlCs characterized experimentally (Profile Dataset worksheet, rhamnose.xlsx), nine have been reported to have 3,5-epimerase activity; these proteins have not been tested for 3-epimerase activity. *Streptomyces niveus* RmlC has both 3-epimerase (major product) and 3,5-epimerase (minor product) activities (95, 96). *Streptomyces bikiniensis* RmlC has only 3-epimerase activity (97). *Streptomyces sp.* GERI-155 RmlC has been characterized as RmlC 3,5-epimerase or 3-epimerase (98). Pairwise sequence identity among the 12 RmlCs varies from 27 to 95%; all alignments are end-to-end. Features that endow only 3,5-epimerase or 3-epimerase activity are not clear. Therefore, a common HMM profile was generated for RmlC.

#### Uger

The Extend dataset is same as the Exp dataset since no new sequence met the criteria required for inclusion in the Extend dataset.

### Searching for homologs of Rmd

The number of Rmds characterized experimentally is just two (15, 99) and hence, instead of generating an HMM profile for this functional family, these two Rmds were used as query to search for homologs by BLASTp (details of these two proteins are given in the Profile Dataset worksheet of rhamnose.xlsx). Gmds from *Pseudomonas aeruginosa* (41), *Aneurinibacillus thermoaerophilus* L420-91T (15) and *Paramecium bursaria Chlorella* virus 1 (42) additionally have weaker (relative to Rmd in the same organism) Rmd activity. This is because both Gmd and Rmd belong to the short chain dehydrogenase/reductase (SDR) protein family, Gmd is the closest paralog of Rmd (43-51% sequence similarity and 67-99% query coverage) and the 3D structure of *A. thermoaerophilus* Rmd is superimposable on that of *P. aeruginosa* Gmd. Therefore, query coverage >=98% and sequence similarity >=90% were used as the key criteria for BLASTp hits. This level of stringency was found to filter out Gmd.

### Fusion proteins with incomplete annotations

In databases, some of the fusion proteins have been assigned the function of only one domain. Such entries get wrongly classified as false negatives when they are obtained as hits for the HMM profile corresponding to the unannotated domain. Two illustrative examples are discussed here to show how such wrong classifications were eliminated. One of the hits to RmlA profile is annotated as RmlB. The length of the hit suggested that this might be a fusion protein. This hit was scored using RmlB profile also. The regions of the hit that align to RmlA and RmlB profiles are distinct suggesting that the hit is a RmlA-RmlB fusion protein and that its annotation in the database is partial. Another hit to the RmlA profile is annotated as WecD (dTDP-fucosamine acetyltransferase). The N-terminal domain of the hit aligns with the RmlA profile. Pairwise sequence alignment (using BLASTp) of the C-terminal domain against known WecDs showed that the C-terminal domain is WecD homolog.

### Analysis of hits from the TrEMBL database and genomes

UniProt uses SAAS and UniRule for electronic annotation of TrEMBL entries. TrEMBL hits were classified by comparing TrEMBL annotation with that of profile annotation (Table 1). Hits for which molecular function is incomplete or not assigned were further classified. For hits whose molecular function does not match with that of the profile annotation, possible reason(s) for the mismatch were analysed by scanning them against HMM profiles / BLASTp corresponding to the annotated activity; for many hits, mismatch could be resolved using this approach. Some hits have biological process, cellular component or incomplete molecular function annotation; some have no annotation whatsoever. Such hits were assigned molecular function by manual curation. For hits which have incomplete molecular function annotation, the profile annotation can be either consistent or inconsistent. For example, an RmlA (a thymidylyltransferase) hit is annotated as nucleotidyltransferase and this is consistent. In contrast, an RmlB/Ugd (a dehydratase) hit is annotated as serine carboxypeptidase and this is inconsistent. In some cases, the inconsistency could be rationalised: an RmlB/Ugd (a dehydratase) hit is annotated as NAD-dependent epimerase, and dehydratases and epimerases which use nucleotide-sugars as substrates belong to SDR superfamily.

### Finding genomic context in genomes

The dataset containing the unique set of proteins from all genomes was scanned against all HMM profiles to identify homologs. These were mapped to their respective genomes (Genome Hits worksheet of rhamnose.xlsx) and their loci were extracted from the feature table of each genome. An extra column was added to the feature table to denote the locus/order of the protein in the genome. The transcripts were numbered sequentially starting from the 5’ end of the genome. The letters “C” and “P” were prefixed to the number to denote the chromosomal and plasmid origin, respectively. In organisms which have more than one plasmid, plasmids were numbered sequentially as “P1.”, “P2.”, etc. If, in a genome, homologs for all enzymes of a pathway are present, then these homologs were classified as follows:

i. Contiguous: the location of the homologs is contiguous in the genome; homologs need not necessarily be in the same strand of DNA.
ii. Neighbourhood: homologs are not contiguous in the genome but are located within 10 genes of each other.
iii. Dispersed: homologs are dispersed in the genome.

In case of (i) and (ii), annotations of neighbouring proteins were extracted to decipher the possible biological function of these homologs. The annotations of the neighbouring genes were categorised as follows:

i. Possibly related – other than transferases: Proteins which might be involved in the same biological process as rhamnose based on literature, i.e., CPS/EPS/LPS/capsid biosynthesis and assembly, cell surface appendage biosynthesis or modification and assembly, secondary metabolite production.
ii. Possibly related – transferases: Proteins which are annotated as transferases, e.g., methyl- and glycosyl-transferases, etc.
iii. Maybe related: Proteins which might be involved in the same biological process as rhamnose, but their substrate specificity is not known. For e.g., epimerases, dehydratases, ABC transporters, etc.
iv. No annotation: Proteins which do not have any annotation or are annotated as DUFs.
v. Not (directly) related: Proteins which are not related to the biological processes which rhamnose is known to be involved in. For e.g., L-histidine biosynthesis, transcription and translation factors, etc.

Proteins in the “Possibly related – other than transferases” category were further sub-grouped as follows: (a) CPS/EPS/LPS/Capsid biosynthesis, (b) secondary metabolite production, (c) cell surface appendage biosynthesis and modification and (d) Non-specific. The “non-specific” category includes genes which might be involved in multiple biological functions such as Qui4N is involved in both LPS biosynthesis and flagellin modification. Based on the number of proteins with these types of annotations present in the neighbourhood of rhamnose biosynthesis genes, additional biological process level annotations were provided to rhamnose biosynthesis genes.

### Identifying homologs of rhamnose biosynthesis genes in metagenomes

The total set of proteins from human-associated and environmental metagenomes was scanned against all HMM profiles to identify homologs of rhamnose biosynthesis genes. Hits were mapped to their respective runs and BioSamples. The BioSamples were mapped to their respective BioProjects and BioProject to their Biomes. The hits found were collated at the level of BioSample.

**ABBREVIATIONS**
HMM: Hidden Markov Model
CPS: Capsular polysaccharide
EPS: Extracellular polysaccharide
LPS: Lipopolysaccharide
CPA: Common polysaccharide antigen
OSA: O-specific antigen

## DECLARATIONS

### Availability of data and materials

All data generated during the analysis of the TrEMBL database, completely sequenced genomes and metagenomes are included in this published article and its supplementary information files.

### Competing interests

All authors declare that they have no competing interests.

### Funding

The fellowship of TM was provided by Department of Science and Technology under the INSPIRE Scheme [INSPIRE code: IF170299].

### Authors’ contributions

PVB conceived and designed the study. TM carried out the work. TM and PVB analysed and interpreted data, and wrote the manuscript. Both authors have read and approved the final manuscript.

## Acknowledgements

TM thanks the Department of Science and Technology, Government of India, for INSPIRE fellowship. The authors thank Ms. Jaya Srivastava and Dr. Shradha Khater for helpful discussion. Authors also thank the Resource for Biocomputing, Visualization, and Informatics at the University of California, San Francisco, which developed the molecular graphics and analyses software UCSF Chimera with support from NIH P41-GM103311. The authors are grateful to various funding agencies and researchers across the world for supporting/making data freely accessible. The authors thank Indian Institute of Technology Bombay for providing facilities and other infrastructure for carrying out this work.

## ADDITIONAL FILES

Fig. S1 Scatter plot of bit score of RmlC hit (against RmlC profile) versus the bit score of RmlD hit (against RmlD profile) from the same genome.

Fig. S2 Bit score distribution of rml hits when they are in contiguity (blue), neighbourhood (red) or dispersed (green) with respect to other rml pathway hits within the same genome.

Fig. S3 Distribution of the number of strains of a species.

Fig. S4 Receiver Operator Characteristics (ROC) curves for functional family-specific HMM profiles RmlA (a), RmlB+Ugd (b), RmlC (c), RmlD (d), Uger (e) and Gmd (f).

Fig. S5 Proteins encoded by *rmlC* may catalyse either 3-epimerization and/or 3,5-epimerization of (d)TDP-4-keto-6-deoxy-glucose.

Table S1 Analysis of hits from the Pfam database

Table S2 Analysis of hits from the CATH database

Table S3 Analysis of composition of biomes at phyla, class, family, genus or species level

Table S4 Databases and software tools used in the present study

Table S5 Functionally important residues in enzymes of rhamnose biosynthesis pathways

Table S6 Number of hits that satisfy the three criteria for hits¶

Excel workbook rhamnose.xlsx: description of training set proteins and data pertaining to (i) validation of profiles, (ii) resolving annotations of TrEMBL and genome hits, (iii) summary of hits obtained from TrEMBL and genomes, (iv) prevalence of the three pathways at domain, phyla, species and strain level, and (v) hits from metagenomes. The “Summary” worksheet has a description of the data in different worksheets.

## REFERENCES

1. Mistou M-Y, Sutcliffe IC, van Sorge NM. Bacterial glycobiology: rhamnose-containing cell wall polysaccharides in Gram-positive bacteria. FEMS Microbiol Rev. 2016;40(4):464–79.

2. Steiner K, Pohlentz G, Dreisewerd K, Berkenkamp S, Messner P, Peter-Katalinić J, et al. New Insights into the Glycosylation of the Surface Layer Protein SgsE from Geobacillus stearothermophilus NRS 2004/3a. Journal of Bacteriology. 2006 Nov 15;188(22):7914–21.

3. Kim BG, Yang SM, Kim SY, Cha MN, Ahn J-H. Biosynthesis and production of glycosylated flavonoids in Escherichia coli: current state and perspectives. Appl Microbiol Biotechnol. 2015 Apr 1;99(7):2979–88.

4. Moses T, Papadopoulou KK, Osbourn A. Metabolic and functional diversity of saponins, biosynthetic intermediates and semi-synthetic derivatives. Crit Rev Biochem Mol Biol. 2014 Nov;49(6):439–62.

5. Slámová K, Kapešová J, Valentová K. “Sweet Flavonoids”: Glycosidase-Catalyzed Modifications. Int J Mol Sci [Internet]. 2018 Jul 21 [cited 2019 Oct 17];19(7). Available from: https://www.ncbi.nlm.nih.gov/pmc/articles/PMC6073497/

6. Boels IC, Beerthuyzen MM, Kosters MHW, Van Kaauwen MPW, Kleerebezem M, De Vos WM. Identification and functional characterization of the Lactococcus lactis rfb operon, required for dTDP-rhamnose Biosynthesis. J Bacteriol. 2004 Mar;186(5):1239–48.

7. Hancock LE, Gilmore MS. The capsular polysaccharide of Enterococcus faecalis and its relationship to other polysaccharides in the cell wall. Proc Natl Acad Sci U S A. 2002 Feb 5;99(3):1574–9.

8. Rigottier-Gois L, Madec C, Navickas A, Matos RC, Akary-Lepage E, Mistou M-Y, et al. The Surface Rhamnopolysaccharide Epa of Enterococcus faecalis Is a Key Determinant of Intestinal Colonization. J Infect Dis. 2015 Jan 1;211(1):62–71.

9. Jiang X-M, Neal B, Santiago F, Lee SJ, Romana LK, Reeves PR. Structure and sequence of the rfb (O antigen) gene cluster of Salmonella serovar typhimurium (strain LT2). Molecular Microbiology. 1991 Mar 1;5(3):695–713.

10. Liu B, Knirel YA, Feng L, Perepelov AV, Senchenkova SN, Wang Q, et al. Structure and genetics of Shigella O antigens. FEMS Microbiol Rev. 2008 Jul;32(4):627–53.

11. Stenutz R, Weintraub A, Widmalm G. The structures of Escherichia coli O-polysaccharide antigens. FEMS Microbiol Rev. 2006 May;30(3):382–403.

12. Wang S, Hao Y, Lam JS, Vlahakis JZ, Szarek WA, Vinnikova A, et al. Biosynthesis of the Common Polysaccharide Antigen of Pseudomonas aeruginosa PAO1: Characterization and Role of GDP-D-Rhamnose:GlcNAc/GalNAc-Diphosphate-Lipid α1,3-D-Rhamnosyltransferase WbpZ. J Bacteriol. 2015 Jun 15;197(12):2012–9.

13. Ovod V, Rudolph K, Knirel Y, Krohn K. Immunochemical characterization of O polysaccharides composing the alpha-D-rhamnose backbone of lipopolysaccharide of Pseudomonas syringae and classification of bacteria into serogroups O1 and O2 with monoclonal antibodies. J Bacteriol. 1996 Nov;178(22):6459–65.

14. Takeuchi K, Ono H, Yoshida M, Ishii T, Katoh E, Taguchi F, et al. Flagellin Glycans from Two Pathovars of Pseudomonas syringae Contain Rhamnose in d and l Configurations in Different Ratios and Modified 4-Amino-4,6-Dideoxyglucose. Journal of Bacteriology. 2007 Oct 1;189(19):6945–56.

15. Kneidinger B, Graninger M, Adam G, Puchberger M, Kosma P, Zayni S, et al. Identification of Two GDP-6-deoxy-d-lyxo-4-hexulose Reductases Synthesizing GDP-d-rhamnose in Aneurinibacillus thermoaerophilus L420-91T. J Biol Chem. 2001 Feb 23;276(8):5577–83.

16. Stephenson AE, Wu H, Novak J, Tomana M, Mintz K, Fives-Taylor P. The Fap1 fimbrial adhesin is a glycoprotein: antibodies specific for the glycan moiety block the adhesion of Streptococcus parasanguis in an in vitro tooth model. Molecular Microbiology. 2002;43(1):147–57.

17. Guerry P, Ewing CP, Schirm M, Lorenzo M, Kelly J, Pattarini D, et al. Changes in flagellin glycosylation affect Campylobacter autoagglutination and virulence. Molecular Microbiology. 2006;60(2):299–311.

18. Takeuchi K, Taguchi F, Inagaki Y, Toyoda K, Shiraishi T, Ichinose Y. Flagellin Glycosylation Island in Pseudomonas syringae pv. glycinea and Its Role in Host Specificity. Journal of Bacteriology. 2003 Nov 15;185(22):6658–65.

19. Ridley BL, O’Neill MA, Mohnen D. Pectins: structure, biosynthesis, and oligogalacturonide-related signaling. Phytochemistry. 2001 Jul 1;57(6):929–67.

20. Lee S-H, Ko C-I, Ahn G, You S, Kim J-S, Heu MS, et al. Molecular characteristics and anti-inflammatory activity of the fucoidan extracted from Ecklonia cava. Carbohydrate Polymers. 2012 Jun 20;89(2):599–606.

21. Rho HS, Ghimeray AK, Yoo DS, Ahn SM, Kwon SS, Lee KH, et al. Kaempferol and Kaempferol Rhamnosides with Depigmenting and Anti-Inflammatory Properties. Molecules. 2011 Apr;16(4):3338–44.

22. Choi HJ, Song JH, Park KS, Kwon DH. Inhibitory effects of quercetin 3-rhamnoside on influenza A virus replication. European Journal of Pharmaceutical Sciences. 2009 Jun 28;37(3):329–33.

23. Choi H-J, Kim J-H, Lee C-H, Ahn Y-J, Song J-H, Baek S-H, et al. Antiviral activity of quercetin 7-rhamnoside against porcine epidemic diarrhea virus. Antiviral Research. 2009 Jan 1;81(1):77–81.

24. Hayder N, Bouhlel I, Skandrani I, Kadri M, Steiman R, Guiraud P, et al. In vitro antioxidant and antigenotoxic potentials of myricetin-3-o-galactoside and myricetin-3-o-rhamnoside from Myrtus communis: Modulation of expression of genes involved in cell defence system using cDNA microarray. Toxicology in Vitro. 2008 Apr 1;22(3):567–81.

25. Tatsimo SJN, Tamokou J de D, Havyarimana L, Csupor D, Forgo P, Hohmann J, et al. Antimicrobial and antioxidant activity of kaempferol rhamnoside derivatives from Bryophyllum pinnatum. BMC Research Notes. 2012 Mar 20;5(1):158.

26. Chen R, Meng F, Liu Z, Chen R, Zhang M. Antitumor activities of different fractions of polysaccharide purified from Ornithogalum caudatum Ait. Carbohydrate Polymers. 2010 May 5;80(3):845–51.

27. Diantini A, Subarnas A, Lestari K, Halimah E, Susilawati Y, Supriyatna S, et al. Kaempferol-3-O-rhamnoside isolated from the leaves of Schima wallichii Korth. inhibits MCF-7 breast cancer cell proliferation through activation of the caspase cascade pathway. Oncology Letters. 2012 May 1;3(5):1069–72.

28. Kim Y-K, Kim YS, Choi SU, Ryu SY. Isolation of flavonol rhamnosides fromloranthus tanakae and cytotoxic effect of them on human tumor cell lines. Arch Pharm Res. 2004 Jan 1;27(1):44–7.

29. Oka T, Nemoto T, Jigami Y. Functional analysis of Arabidopsis thaliana RHM2/MUM4, a multidomain protein involved in UDP-D-glucose to UDP-L-rhamnose conversion. J Biol Chem. 2007 Feb 23;282(8):5389–403.

30. Xu Y, Singh KV, Qin X, Murray BE, Weinstock GM. Analysis of a Gene Cluster of Enterococcus faecalis Involved in Polysaccharide Biosynthesis. Infect Immun. 2000 Feb;68(2):815–23.

31. Rahim R, Burrows LL, Monteiro MA, Perry MB, Lam JS. Involvement of the rml locus in core oligosaccharide and O polysaccharide assembly in Pseudomonas aeruginosa. Microbiology (Reading, Engl). 2000 Nov;146 (Pt 11):2803–14.

32. Chiang SL, Mekalanos JJ. rfb mutations in Vibrio cholerae do not affect surface production of toxin-coregulated pili but still inhibit intestinal colonization. Infect Immun. 1999 Feb;67(2):976–80.

33. Yamashita Y, Tomihisa K, Nakano Y, Shimazaki Y, Oho T, Koga T. Recombination between gtfB and gtfC is required for survival of a dTDP-rhamnose synthesis-deficient mutant of Streptococcus mutans in the presence of sucrose. Infect Immun. 1999 Jul;67(7):3693–7.

34. Burns SM, Hull SI. Comparison of loss of serum resistance by defined lipopolysaccharide mutants and an acapsular mutant of uropathogenic Escherichia coli O75:K5. Infection and immunity. 1998;66(9):4244–53.

35. Kantardjieff KA, Kim C-Y, Naranjo C, Waldo GS, Lekin T, Segelke BW, et al. Mycobacterium tuberculosis RmlC epimerase (Rv3465): a promising drug-target structure in the rhamnose pathway. Acta Cryst D. 2004 May 1;60(5):895–902.

36. Ma Y, Pan F, McNeil M. Formation of dTDP-Rhamnose Is Essential for Growth of Mycobacteria. Journal of Bacteriology. 2002 Jun 15;184(12):3392–5.

37. Ma Y, Stern RJ, Scherman MS, Vissa VD, Yan W, Jones VC, et al. Drug Targeting Mycobacterium tuberculosis Cell Wall Synthesis: Genetics of dTDP-Rhamnose Synthetic Enzymes and Development of a Microtiter Plate-Based Screen for Inhibitors of Conversion of dTDP-Glucose to dTDP-Rhamnose. Antimicrobial Agents and Chemotherapy. 2001 May 1;45(5):1407–16.

38. Ashburner M, Ball CA, Blake JA, Botstein D, Butler H, Cherry JM, et al. Gene ontology: tool for the unification of biology. The Gene Ontology Consortium. Nat Genet. 2000 May;25(1):25–9.

39. Aguirrezabalaga I, Olano C, Allende N, Rodriguez L, Braña AF, Méndez C, et al. Identification and expression of genes involved in biosynthesis of L-oleandrose and its intermediate L-olivose in the oleandomycin producer Streptomyces antibioticus. Antimicrob Agents Chemother. 2000 May;44(5):1266–75.

40. El-Gebali S, Mistry J, Bateman A, Eddy SR, Luciani A, Potter SC, et al. The Pfam protein families database in 2019. Nucleic Acids Res. 2019 Jan 8;47(D1):D427–32.

41. King JD, Poon KKH, Webb NA, Anderson EM, McNally DJ, Brisson J-R, et al. The structural basis for catalytic function of GMD and RMD, two closely related enzymes from the GDP-d-rhamnose biosynthesis pathway. FEBS J. 2009 May;276(10):2686–700.

42. Tonetti M, Zanardi D, Gurnon JR, Fruscione F, Armirotti A, Damonte G, et al. Paramecium bursaria Chlorella Virus 1 Encodes Two Enzymes Involved in the Biosynthesis of GDP-L-fucose and GDP-D-rhamnose. J Biol Chem. 2003 Jun 13;278(24):21559–65.

43. Kneidinger B, Marolda C, Graninger M, Zamyatina A, McArthur F, Kosma P, et al. Biosynthesis Pathway of ADP-l-glycero-β-d-manno-Heptose in Escherichia coli. J Bacteriol. 2002 Jan;184(2):363–9.

44. Erbel PJA, Barr K, Gao N, Gerwig GJ, Rick PD, Gardner KH. Identification and biosynthesis of cyclic enterobacterial common antigen in Escherichia coli. J Bacteriol. 2003 Mar;185(6):1995–2004.

45. Lam JS, Taylor VL, Islam ST, Hao Y, Kocíncová D. Genetic and Functional Diversity of Pseudomonas aeruginosa Lipopolysaccharide. Front Microbiol [Internet]. 2011 [cited 2019 Sep 4];2. Available from: https://www.frontiersin.org/articles/10.3389/fmicb.2011.00118/full

46. McCaughey LC, Grinter R, Josts I, Roszak AW, Waløen KI, Cogdell RJ, et al. Lectin-Like Bacteriocins from Pseudomonas spp. Utilise D-Rhamnose Containing Lipopolysaccharide as a Cellular Receptor. PLoS Pathog [Internet]. 2014 Feb 6 [cited 2019 Sep 4];10(2). Available from: https://www.ncbi.nlm.nih.gov/pmc/articles/PMC3916391/

47. King JD, Kocíncová D, Westman EL, Lam JS. Review: Lipopolysaccharide biosynthesis in Pseudomonas aeruginosa. Innate Immun. 2009 Oct;15(5):261–312.

48. Bryan BA, Linhardt RJ, Daniels L. Variation in composition and yield of exopolysaccharides produced by Klebsiella sp. strain K32 and Acinetobacter calcoaceticus BD4. Applied and Environmental Microbiology. 1986 Jun;51(6):1304.

49. Kaplan N, Rosenberg E, Jann B, Jann K. Structural studies of the capsular polysaccharide of Acinetobacter calcoaceticus BD4. European Journal of Biochemistry. 1985;152(2):453–8.

50. Morin A, Duchiron F, Monsan PF. Production and recovery of rhamnose-containing polysaccharides from acinetobacter calcoaceticus. Journal of Biotechnology. 1987 Nov 1;6(4):293–306.

51. Rosenberg E, Kaplan N, Pines O, Rosenberg M, Gutnick D. Capsular polysaccharides interfere with adherence of Acinetobacter calcoaceticus to hydrocarbon. FEMS Microbiology Letters. 1983 Mar;17(1–3):157–60.

52. Haseley SR, Wilkinson SG. Structure of the putative O10 antigen from Acinetobacter baumannii. Carbohydrate Research. 1994 Nov 1;264(1):73–81.

53. Haseley SR, Wilkinson SG. Structure of the O-7 antigen from Acinetobacter baumannii. Carbohydrate Research. 1998 Jan 1;306(1):257–63.

54. Vinogradov EV, Petersen BO, Thomas-Oates JE, Duus JØ, Brade H, Holst O. Characterization of a Novel Branched Tetrasaccharide of 3-Deoxy-d-manno-oct-2-ulopyranosonic Acid THE STRUCTURE OF THE CARBOHYDRATE BACKBONE OF THE LIPOPOLYSACCHARIDE FROM ACINETOBACTER BAUMANNII STRAIN NCTC 10303 (ATCC 17904). J Biol Chem. 1998 Oct 23;273(43):28122–31.

55. Haseley SR, Galbraith L, Wilkinson SG. Structure of a surface polysaccharide from Acinetobacter baumannii strain 214. Carbohydrate Research. 1994 May 20;258:199–206.

56. Vinogradov E, Pantophlet R, Dijkshoorn L, Brade L, Holst O, Brade H. Structural and Serological Characterisation of Two O-Specific Polysaccharides of Acinetobacter. European Journal of Biochemistry. 2004 Jul 23;239:602–10.

57. Vinogradov EV, Brade L, Brade H, Holst O. Structural and serological characterisation of the O-antigenic polysaccharide of the lipopolysaccharide from Acinetobacter baumannii strain 24. Carbohydrate Research. 2003 Nov 14;338(23):2751–6.

58. Russo TA, Luke NR, Beanan JM, Olson R, Sauberan SL, MacDonald U, et al. The K1 Capsular Polysaccharide of Acinetobacter baumannii Strain 307-0294 Is a Major Virulence Factor. Infection and Immunity. 2010 Sep;78(9):3993.

59. Jofré E, Lagares A, Mori G. Disruption of dTDP-rhamnose biosynthesis modifies lipopolysaccharide core, exopolysaccharide production, and root colonization in Azospirillum brasilense. FEMS Microbiol Lett. 2004 Feb 1;231(2):267–75.

60. Kaplan JB, Perry MB, MacLean LL, Furgang D, Wilson ME, Fine DH. Structural and Genetic Analyses of O Polysaccharide from Actinobacillus actinomycetemcomitans Serotype f. Infection and Immunity. 2001 Sep;69(9):5375.

61. Raja M, Ummer F, Dhivakar CP. Aggregatibacter Actinomycetemcomitans – A Tooth Killer? Journal of Clinical and Diagnostic Research : JCDR. 2014 Aug;8(8):ZE13.

62. Chen C, Kittichotirat W, Chen W, Downey JS, Si Y, Bumgarner R. Genome Sequence of Naturally Competent Aggregatibacter actinomycetemcomitans Serotype a Strain D7S-1. Journal of Bacteriology. 2010 May 15;192(10):2643–4.

63. Narayanan AM, Ramsey MM, Stacy A, Whiteley M. Defining Genetic Fitness Determinants and Creating Genomic Resources for an Oral Pathogen. Appl Environ Microbiol. 2017 Jul 15;83(14):e00797–17.

64. Vinogradov E, Frirdich E, MacLean LL, Perry MB, Petersen BO, Duus JØ, et al. Structures of lipopolysaccharides from Klebsiella pneumoniae. Eluicidation of the structure of the linkage region between core and polysaccharide O chain and identification of the residues at the non-reducing termini of the O chains. J Biol Chem. 2002 Jul 12;277(28):25070–81.

65. Bebault GM, G.S. Dutton G, Funnell NA, Mackie KL. Structural investigation of Klebsiella serotype K32 polysaccharide. Carbohydrate Research. 1978 Jun 1;63:183–92.

66. Björndal H, Lindberg B, Lönngren J, Rosell KG, Nimmich W. Structural studies of the Klebsiella type 47 capsular polysaccharide. Carbohydr Res. 1973 Apr;27(2):373–8.

67. Choy Y-M, Dutton GGS. The Structure of the Capsular Polysaccharide of Klebsiella K-type 72; Occurrence of 3,4-O-(1-Carboxyethylidene)-L-rhamnose. Can J Chem. 1974 Feb 15;52(4):684–7.

68. Follador R, Heinz E, Wyres KL, Ellington MJ, Kowarik M, Holt KE, et al. The diversity of Klebsiella pneumoniae surface polysaccharides. Microb Genom [Internet]. 2016 Aug 25 [cited 2019 Sep 8];2(8). Available from: https://www.ncbi.nlm.nih.gov/pmc/articles/PMC5320592/

69. Lindberg B, Lönngren J, Thompson JL. Structural studies of the Klebsiella type 9 capsular polysaccharide. Carbohydr Res. 1972 Nov;25(1):49–57.

70. Shu H-Y, Fung C-P, Liu Y-M, Wu K-M, Chen Y-T, Li L-H, et al. Genetic diversity of capsular polysaccharide biosynthesis in Klebsiella pneumoniae clinical isolates. Microbiology (Reading, Engl). 2009 Dec;155(Pt 12):4170–83.

71. Stenutz R, Erbing B, Widmalm G, Jansson PE, Nimmich W. The structure of the capsular polysaccharide from Klebsiella type 52, using the computerised approach CASPER and NMR spectroscopy. Carbohydr Res. 1997 Jul 11;302(1–2):79–84.

72. Yother J. Capsules of Streptococcus pneumoniae and other bacteria: paradigms for polysaccharide biosynthesis and regulation. Annu Rev Microbiol. 2011;65:563–81.

73. Tsukioka Y, Yamashita Y, Nakano Y, Oho T, Koga T. Identification of a fourth gene involved in dTDP-rhamnose synthesis in Streptococcus mutans. J Bacteriol. 1997 Jul;179(13):4411–4.

74. Tsukioka Y, Yamashita Y, Oho T, Nakano Y, Koga T. Biological function of the dTDP-rhamnose synthesis pathway in Streptococcus mutans. J Bacteriol. 1997 Feb;179(4):1126–34.

75. García E, Llull D, Muñoz R, Mollerach M, López R. Current trends in capsular polysaccharide biosynthesis of Streptococcus pneumoniae. Research in Microbiology. 2000 Jul;151(6):429–35.

76. Jiang S-M, Wang L, Reeves PR. Molecular Characterization of Streptococcus pneumoniae Type 4, 6B, 8, and 18C Capsular Polysaccharide Gene Clusters. Infect Immun. 2001 Mar;69(3):1244–55.

77. Morona JK, Morona R, Paton JC. Comparative genetics of capsular polysaccharide biosynthesis in Streptococcus pneumoniae types belonging to serogroup 19. J Bacteriol. 1999 Sep;181(17):5355–64.

78. Morona JK, Morona R, Paton JC. Characterization of the locus encoding the Streptococcus pneumoniae type 19F capsular polysaccharide biosynthetic pathway. Molecular Microbiology. 1997;23(4):751–63.

79. Brennan PJ. Structure, function, and biogenesis of the cell wall of Mycobacterium tuberculosis. Tuberculosis. 2003 Feb 1;83(1):91–7.

80. McNeil M, Daffe M, Brennan PJ. Evidence for the nature of the link between the arabinogalactan and peptidoglycan of mycobacterial cell walls. J Biol Chem. 1990 Oct 25;265(30):18200–6.

81. Babaoglu K, Page MA, Jones VC, McNeil MR, Dong C, Naismith JH, et al. Novel inhibitors of an emerging target in Mycobacterium tuberculosis; substituted thiazolidinones as inhibitors of dTDP-rhamnose synthesis. Bioorganic & Medicinal Chemistry Letters. 2003 Oct 16;13(19):3227–30.

82. Narayanan S, Deshpande U. Whole-Genome Sequences of Four Clinical Isolates of Mycobacterium tuberculosis from Tamil Nadu, South India. Genome Announc. 2013 Jun 20;1(3).

83. Ansaruzzaman M, Albert MJ, Holme T, Jansson P-E, Rahman MM, Widmalm G. A Klebsiella pneumoniae Strain that Shares a Type-Specific Antigen with Shigella flexneri Serotype 6. European Journal of Biochemistry. 1996 May 1;237(3):786–91.

84. Simmons DA. Immunochemistry of Shigella flexneri O-antigens: a study of structural and genetic aspects of the biosynthesis of cell-surface antigens. Bacteriol Rev. 1971 Jun;35(2):117–48.

85. Iguchi A, Iyoda S, Kikuchi T, Ogura Y, Katsura K, Ohnishi M, et al. A complete view of the genetic diversity of the Escherichia coli O-antigen biosynthesis gene cluster. DNA Res. 2015 Feb;22(1):101–7.

86. Orskov I, Orskov F, Jann B, Jann K. Serology, chemistry, and genetics of O and K antigens of Escherichia coli. Bacteriol Rev. 1977 Sep;41(3):667–710.

87. Silver RP, Aaronson W, Vann WF. The K1 capsular polysaccharide of Escherichia coli. Rev Infect Dis. 1988 Aug;10 Suppl 2:S282–286.

88. Whitfield C. Biosynthesis and assembly of capsular polysaccharides in Escherichia coli. Annu Rev Biochem. 2006;75:39–68.

89. Joiner KA. Complement Evasion by Bacteria and Parasites. Annual Review of Microbiology. 1988;42(1):201–30.

90. Wahl HP, Grisebach H. Biosynthesis of streptomycin dTDP-Dihydrostreptose synthase from Streptomyces griseus and dTDP-4-keto-l-rhamnose 3,5-epimerase from S. griseus and Escherichia coli Y10. Biochimica et Biophysica Acta (BBA) – Enzymology. 1979 May 10;568(1):243–52.

91. Mathesius U. Flavonoid Functions in Plants and Their Interactions with Other Organisms. Plants (Basel) [Internet]. 2018 Apr 3 [cited 2019 Oct 17];7(2). Available from: https://www.ncbi.nlm.nih.gov/pmc/articles/PMC6027123/

92. Matsunaga T, Ishii T, Matsumoto S, Higuchi M, Darvill A, Albersheim P, et al. Occurrence of the Primary Cell Wall Polysaccharide Rhamnogalacturonan II in Pteridophytes, Lycophytes, and Bryophytes. Implications for the Evolution of Vascular Plants. Plant Physiol. 2004 Jan;134(1):339–51.

93. Parakkottil Chothi M, Duncan GA, Armirotti A, Abergel C, Gurnon JR, Van Etten JL, et al. Identification of an L-rhamnose synthetic pathway in two nucleocytoplasmic large DNA viruses. J Virol. 2010 Sep;84(17):8829–38.

94. The UniProt Consortium. UniProt: a worldwide hub of protein knowledge. Nucleic Acids Research. 2019 Jan 8;47(D1):D506–15.

95. Tello M, Jakimowicz P, Errey JC, Meyers CLF, Walsh CT, Buttner MJ, et al. Characterisation of Streptomyces spheroides NovW and revision of its functional assignment to a dTDP-6-deoxy-D-xylo-4-hexulose 3-epimerase. Chem Commun. 2006 Mar 1;(10):1079–81.

96. Thu Thuy TT, Lee HC, Kim C-G, Heide L, Sohng JK. Functional characterizations of novWUS involved in novobiocin biosynthesis from Streptomyces spheroides. Archives of Biochemistry and Biophysics. 2005 Apr 1;436(1):161–7.

97. Kubiak RL, Phillips RK, Zmudka MW, Ahn MR, Maka EM, Pyeatt GL, et al. Structural and Functional Studies on a 3′-Epimerase Involved in the Biosynthesis of dTDP-6-deoxy-d-allose. Biochemistry. 2012 Nov 20;51(46):9375–83.

98. Sohng J-K, Kim H, Nam D-H, Lim D-O, Han J-M, Lee H-J, et al. Cloning, expression, and biological function of a dTDP-deoxyglucose epimerase (gerF) gene from Streptomyces sp. GERI-155. Biotechnol Lett. 2004 Feb;26(3):185–91.

99. Mäki M, Järvinen N, Räbinä J, Roos C, Maaheimo H, Renkonen R. Functional expression of Pseudomonas aeruginosa GDP-4-keto-6-deoxy-d-mannose reductase which synthesizes GDP-rhamnose. European Journal of Biochemistry. 2002;269(2):593–601.

